# Histone deacetylase 6 inhibition promotes microtubule acetylation and facilitates autophagosome-lysosome fusion in dystrophin-deficient *mdx* mice

**DOI:** 10.1101/2022.04.29.490072

**Authors:** Akanksha Agrawal, Erin L. Clayton, Courtney L. Cavazos, Benjamin A. Clayton, George G. Rodney

## Abstract

Duchenne Muscular Dystrophy (DMD) is a severe X-linked genetic disorder. Defective autophagy and disorganized microtubule network contributes to DMD pathogenesis, yet the mechanisms by which microtubule alterations regulate autophagy remain elusive. We show decreased acetylated α-tubulin and enhanced histone deacetylase (HDAC6) expression in *mdx* mice. Pharmacological inhibition of HDAC6 increases tubulin acetylation and enhances Q-SNARE complex formation, leading to improved autophagosome-lysosome fusion. HDAC6 inhibition reduces apoptosis, inflammation, muscle damage and prevents contraction induced force loss. HDAC6 inhibition restores peroxiredoxin (PrxII) by increasing its acetylation and protecting it from hyper-oxidation, hence modulating intracellular redox status in *mdx* mice. Genetic inhibition of Nox2 activity in *mdx* mice promotes autophagosome maturation. Our data highlight that autophagy is differentially regulated by redox and acetylation in *mdx* mice. By restoring tubulin acetylation HDAC6 inhibition enhances autophagy, ameliorates the dystrophic phenotype and improves muscle function, suggesting a potential therapeutic target for treating DMD.

## Introduction

Duchenne Muscular Dystrophy (DMD) is the X-linked recessive genetic disorder caused by mutations in *DMD* gene which encodes for dystrophin (Dp427m), a key component of the dystrophin-glycoprotein complex (DGC). It affects approximately 1:4000 to 1:5000 live male births worldwide ^1^ resulting in progressive muscle wasting and degeneration, leading to death due to cardiac dysfunction and respiratory failure ^2^. Over the past two decades, several therapeutic approaches have been evaluated to combat the pathogenesis of the disease, but DMD is still incurable. Novel genetic approaches hold promising therapy; ^3^ however, many challenges still exist due to the variability in exon skipping efficiency among patients, the non-homogenous restoration of dystrophin between muscle types (including absence of effect of current exon skipping agents on cardiac muscle), and minimal alterations in immune cell infiltration. In order to combat these limitations, the treatment plan for DMD will likely entail a combination of genetic and pharmacological interventions.

Dystrophin is a large cytoskeletal protein located at the sarcolemma that mechanically links the internal cytoskeleton to the extracellular matrix and is critical for muscle-membrane stability during contraction ^4^. Lack of dystrophin results in disassembly of the DGC and increases the sarcolemma susceptibility to contraction-induced injury ^5^. This leads to a series of pathological events including, increased ROS signaling, aberrant Ca^2+^ release, inflammation, impaired autophagy, fibrosis, apoptosis, and decreased force production. ^5–7^

Macroautophagy, hereafter referred to as autophagy, is a highly conserved process involving a series of sequential events for bulk degradation of cytosolic components and organelles through delivery of autophagosomes to lysosomes, and thus, maintaining cellular homeostasis.^8^ Accumulating evidence shows that impaired autophagy contributes to muscle weakness and cell death in both *mdx* mice (mouse model of DMD) and DMD patients ^9, 10^. Recent work from our lab has shown that Nox2/Src kinase impairs autophagy by regulating the PI3K/Akt/mTOR pathway in *mdx* mice ^7^. To better comprehend how defective autophagy leads to DMD pathophysiology, and to develop therapeutic strategies, we need to elucidate the defects at different steps of the autophagic pathway.

Dystrophin is a microtubule-associated protein found to bind microtubules (MT). ^11^ The MT lattice becomes disorganized when dystrophin expression is ablated as in the *mdx* mouse ^11–13^. We have shown that the increased Nox2-ROS observed in *mdx* skeletal muscle regulates the MT network^14^. MTs undergo post-translational modifications that regulate their biological functions. Some studies suggest detyrosination of α-tubulin increases muscle stiffness and decreases force production in *mdx* mice ^15, 16^. In non-muscle cells, acetylation of α-tubulin has been shown to regulate the formation of pre-autophagosomal structures, vesicular movements and autophagosome-lysosome fusion ^17–19^. Acetylated MTs recruit the motor protein kinesin-1 to transport autophagosomes in a cargo-specific manner along the MT tracks. Changes in MT acetylation leads to alterations in MT dynamics and organization, cell migration, and autophagy^18, 20^. Despite the extensive investigations largely based on MT dynamics, stability and its altered network in dystrophic mice, no study has addressed the role of acetylated MTs specifically on autophagosome biogenesis and autophagosome-lysosome fusion in DMD. Therefore, the present study was designed to unravel the plausible mechanisms underpinning the role of MTs in regulating autophagy in dystrophic mice.

Histone deacetylases (HDACs) are a class of deacetylase enzymes involved in chromatin remodeling and gene expression ^21^. Currently, epigenetic drugs (e.g. Givinostat) targeting HDACs are in phase III clinical trials to assess their functional effects in DMD patients ^2, 22^. Recently, HDAC6 (Class IIb) inhibition has emerged as one potential selective pharmacological target in neurodegenerative diseases. HDAC6 catalyzes the deacetylation of non-histone proteins such as α-tubulin, leading to altered MT stability and organization ^23^. In addition, HDAC6 can control redox regulation through acetylation of peroxiredoxin (Prx1 and PrxII)^24, 25^. Therefore, selective HDAC6 inhibition has the potential to reduce the toxicity related to the off-target effects of pan-HDAC inhibitors (Givinostat). ^26^ Recent investigations have suggested that the HDAC6 inhibitor, Tubastatin A (TubA) promotes MT acetylation, improving autophagic flux, redox balance, and functional recovery in neurodegenerative disorders ^19, 27–29^, cardiomyopathy ^30^, idiopathic pulmonary fibrosis^31^, myocardial ischemia/reperfusion injury ^32^, osteoarthritis ^33^, and kidney injury ^34^. In the current study, we show differential regulation of autophagy in *mdx* skeletal muscle; while autophagosome maturation is regulated by Nox2-ROS, autophagosome-lysosome fusion is regulated by MT acetylation. Furthermore, we discovered the therapeutic efficacy of HDAC6 inhibitor, TubA, in promoting MT acetylation and restoring autophagic flux in *mdx* mice. TubA treatment significantly restricted muscle damage and apoptosis and induced muscle functional recovery.

## Results

### Impaired autophagosomal biogenesis/maturation in *mdx* mice

The maturation of double-membrane vesicle structures called autophagosomes occurs in a highly orchestrated manner to achieve successful delivery to the lysosomes for fusion ^35^. One of the initial steps in autophagosome biogenesis is the recruitment and activation of the class III phosphatidylinositol 3-kinase complex (PI3K), consisting of Beclin, ATG14L, VPS34, and VPS15 to facilitate the phagophore nucleation^36^. Immunoblot analyses of TA muscle homogenate revealed a significant decrease in ATG14L and VPS34, whereas both Beclin and VPS15 were found to be increased in *mdx* muscles as compared to WT **(Fig.1a-b).** Immunoprecipitation of Beclin followed by western blot for ATG14L showed decreased ATG14L/Beclin complex formation in *mdx* muscle as compared to WT while Bcl-2/Beclin complex formation was not altered **(Fig. 1c).** These data are consistent with inhibition of vesicle nucleation in *mdx* mice. Autophagy induction by heterodimerization of Beclin with ATG14L-VPS34 is regulated by activated JNK. JNK-interacting protein 1 (JIP-1) binds protein kinases (e.g. MAPKK/MKK7) to promote phosphorylation and activation of JNK ^37^. We observed that the protein expressions levels of JIP- 1 and p-JNK (Thr183/Ty185) are downregulated in *mdx* skeletal muscle, whereas MAPKK was significantly upregulated in *mdx* mice **(Fig 1d-e).** P-JNK phosphorylates Bcl-2 (S70), which disrupts the Bcl-2/Beclin complex, allowing Beclin to bind the PI3KclassIII complex for phagophore nucleation. We found lower levels of p-Bcl-2 in *mdx* muscles as compared to WT, likely due to the decrease in p-JNK levels **(Fig 1d-e).** Together, these observations suggest that phagophore nucleation is disrupted due to inhibition of JIP-1/JNK activation and subsequent phosphorylation of Bcl-2 in *mdx* mice.

**Fig. 1.**
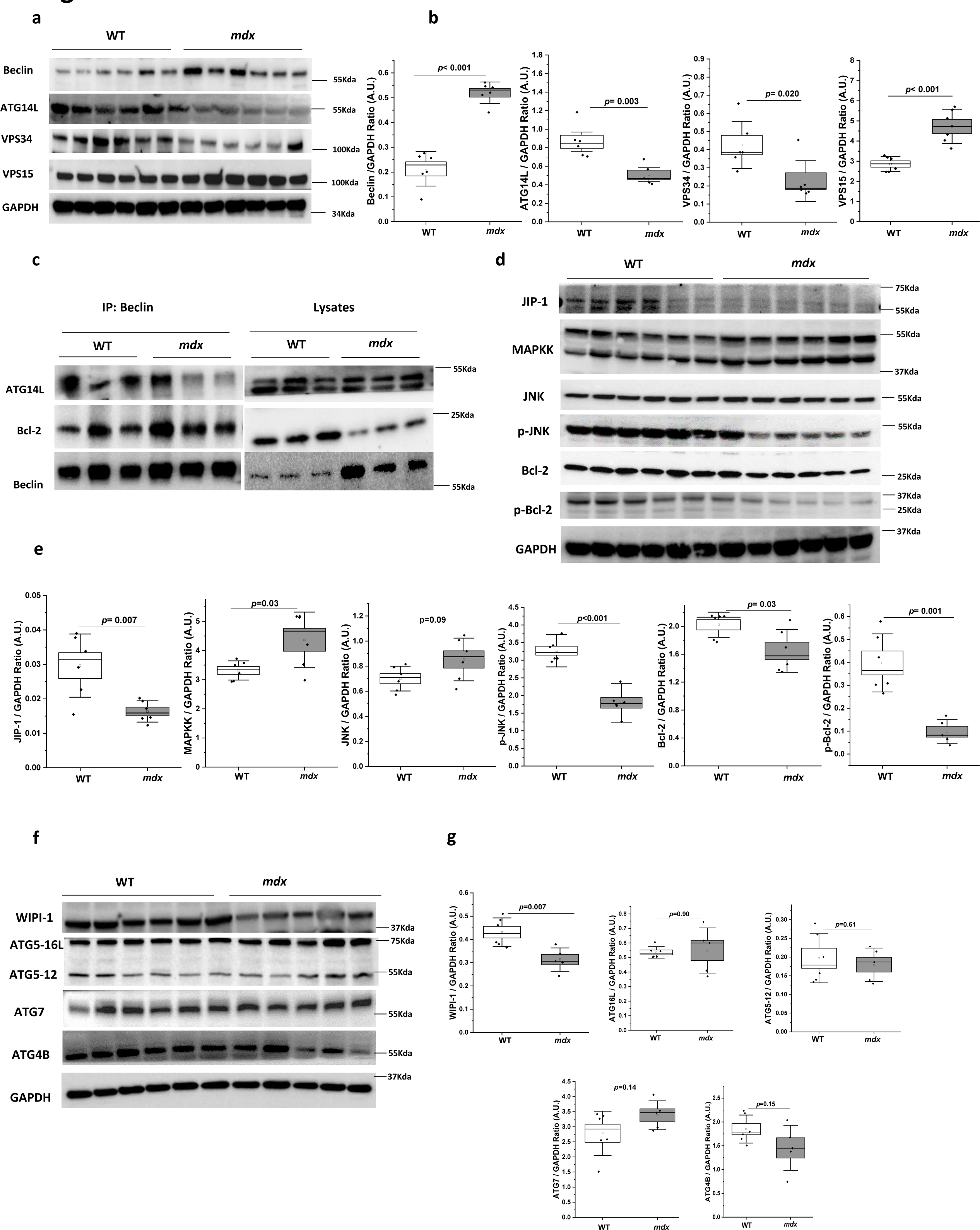
Impaired autophagosomal biogenesis/maturation in *mdx* mice. **(a-b)** Immunoblot and densitometry analysis for autophagy regulatory proteins involved in vesicle nucleation (n=6 per group) and (ATG14L; WT, n = 6; *mdx*, n = 5). **(c)** Skeletal muscle tissue lysate prepared and subjected to immunoprecipitation using anti-Beclin, and the associated ATG14L, and Bcl-2 were determined using immunoblotting (n=3 per group). **(d-e)** JIP-1/JNK signaling was detected by immunoblot with antibodies as indicated (n = 6 per group) and (p-Bcl-2; WT, n=6; *mdx*,n=5). Densitometry analysis of immunoblots represented by box plots**. (f-g)** Immunoblot for proteins involved in autophagosome maturation and elongation (WT, n = 6; *mdx*, n = 5). Densitometry analysis of immunoblot represented by graph. GAPDH served as a loading control. Statistical difference between two groups were determined using two-sample student-*t-*test and Welch’s correction test was performed for non-equal variance between two groups.

Vesicle nucleation is followed by elongation and expansion of the phagophore in the cytoplasm, which is regulated by the ATG5-12 complex. Our data revealed that WIPI-1 (WD Repeat Domain Phosphoinositide Interacting 1), an early marker of autophagosome formation which fosters the recruitment of downstream ATGs, was significantly downregulated in *mdx* muscle **(Fig. 1f-g)** whereas ATG7, ATG4B, and ATG5-12 did not show any prominent change in *mdx* as compared to WT **(Fig. 1f-g).** In addition, gene expression analysis of autophagy related markers *uvrag, vps34, atg14l, FIP200, atg5, atg12, and gabarapl1* did not show any change in *mdx* skeletal muscle as compared to WT **(Fig.S1**). Together, our data suggests that defects in the autophagosome maturation is due to inhibition of vesicle nucleation and elongation of the phagophore in *mdx* skeletal muscle.

### Impairment in the SNARE-mediated autophagosomal-lysosomal fusion in *mdx* mice

To clear sequestered cytosolic components the autophagosome must fuse with the lysosome, forming the autolysosome. The process of autophagosome-lysosome fusion is regulated by the SNARE tertiary complex STX17-SNAP29-VAMP8. To achieve autophagosome-lysosome fusion, autophagosome-localized Q-SNARE STX17 must interact with SNAP29 and the lysosomal-localized R-SNARE VAMP8. ATG14L acts as a tethering factor to facilitate the fusion by directly interacting with the STX17-SNAP29 binary complex, and primes it for binding to VAMP8 on lysosomes ^38^ . We found that STX17 was significantly decreased in *mdx* as compared to WT, while VAMP8 and SNAP29 both showed no change **(Fig.2a-b).** CO-IP with anti-SNAP29 in TA muscle lysates revealed reduced interaction of SNAP29 with STX17 and VAMP8 in *mdx* muscle whereas SNAP29 retained binding with ATG14L **(Fig. 2c)**. CO-IP with an anti-VAMP8 showed less interaction of VAMP8 with STX17, SNAP29, and ATG14L **(Fig. S2a),** confirming reduced autophagosome-lysosome fusion in *mdx* skeletal muscle.

**Fig. 2.**
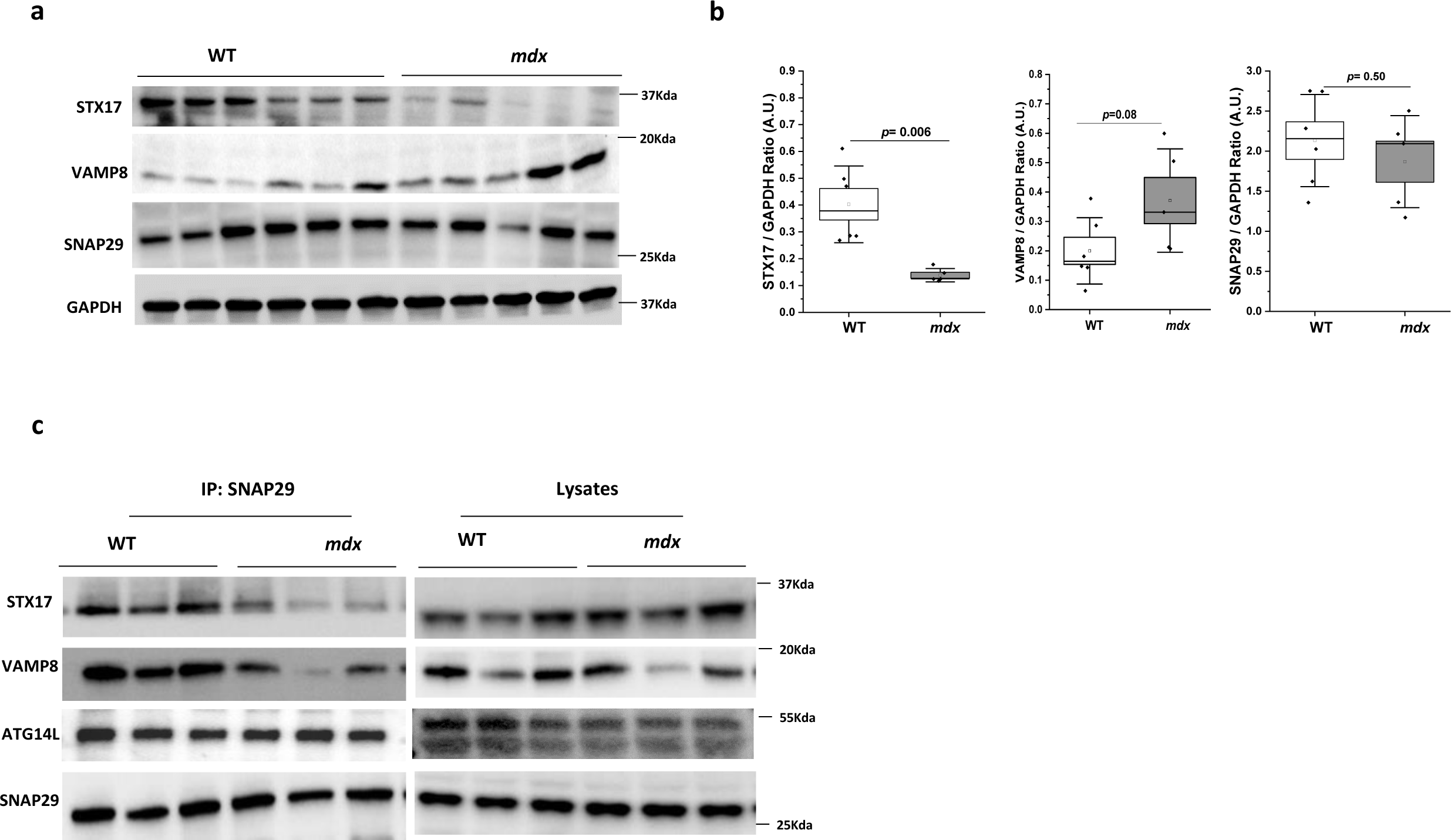
Impairment in the SNARE complex-mediated autophagosomal-lysosomal fusion in *mdx* mice. **(a-b)** Protein expressions of SNARE tertiary complex-STX17, VAMP8, and SNAP29 was determined by immunoblot (WT, n = 6; *mdx*, n = 5). Densitometry analysis of immunoblot represented by box plots. **(c)** Skeletal muscle lysate was prepared and subjected to immunoprecipitation using anti-SNAP29, and the associated SNAP29, STX17, ATG14L, and VAMP8 were determined using immunoblotting (n=3 per group). GAPDH served as a loading control. Statistical difference between two groups were determined using two-sample student-*t-*test and Welch’s correction test was performed for non-equal variance between two groups.

Acetylation of STX17 inhibits the interaction between STX17 and SNAP29, and formation of the Q-SNARE complex^39^. Therefore, we asked whether STX17 acetylation is impaired in *mdx* muscle, leading to reduced interaction between the Q-SNARE complexes. We observed decreased acetylation of STX17 in *mdx* muscle compared to WT. We also found increased expression of HDAC2, the primary deacetylase of STX17 ^39^ in *mdx* muscle **(Fig. S2b-d)**. These data suggest that the reduced interaction in the Q-SNARE complex in *mdx* muscle is not due to increased acetylation of STX17.

### Genetic ablation of p47^phox^ promotes autophagosomal maturation without facilitating autolysosome formation in *mdx* mice

We have previously shown that genetic deletion of p47^phox^ function in *mdx* mice protects against oxidative stress and improves autophagy (i.e. increased LC3II/I and decreased p62) compared with *mdx* ^7^ . Having established defects at multiple stages in autophagy, including autophagosome maturation and autolysosome formation in *mdx* mice, we asked whether these steps are regulated by Nox2-ROS in *mdx* skeletal muscle. We found increased p-JNK, p-Bcl-2 and JIP-1 protein levels (**Fig. 3b-c**), as well as increased Beclin-ATG14L complex formation and a decrease in the Beclin- Bcl-2 complex (**Fig. 3a**); all consistent with improved vesicle nucleation. We also found an increase in the autophagy elongation complex, ATG5-12 in p47^-/-^/*mdx* mice **(Fig. 3d-e).** We then determined whether autophagosome-lysosome fusion is affected by genetic deletion of p47^phox^ in *mdx* mice. Interestingly, we did not find any improvement in the interaction between STX17- SNAP29-VAMP8 tertiary complexes in p47^-/-^/*mdx* mice as compared to *mdx* mice **(Fig 3f).** In addition, no changes were found in the lysosomal protein LAMP2 and lysosomal hydrolase, Cathepsin B in p47^-/-^/*mdx* mice as compared with *mdx* (**Fig. 3g-h**). Overall, our data indicates that inhibition of Nox2-ROS promotes autophagosome maturation but does not enhance the fusion of autophagosomes with lysosomes in *mdx* mice.

**Fig. 3.**
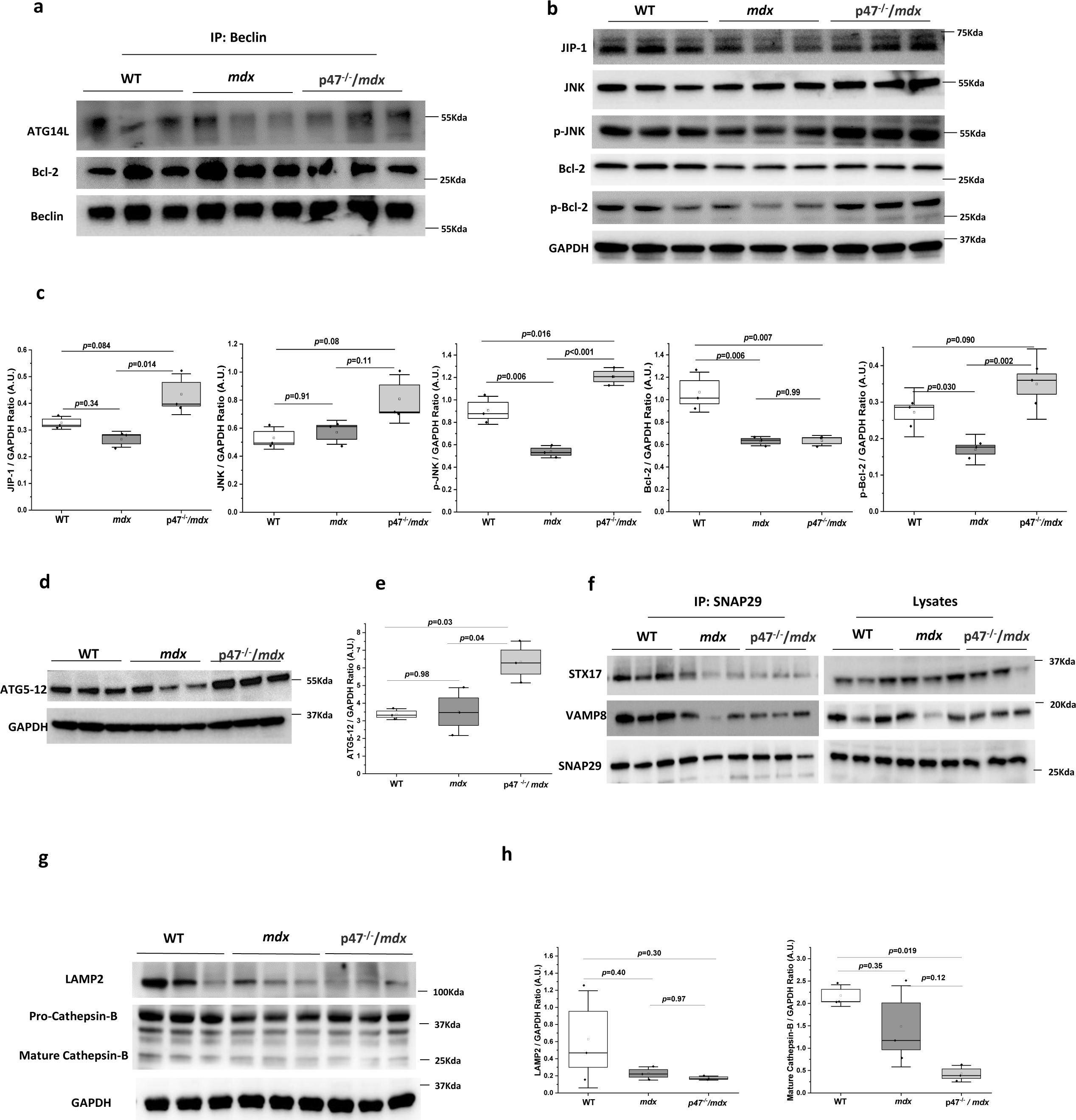
Genetic ablation of p47^phox^ promotes autophagosomal maturation in *mdx* mice. **(a)** Skeletal muscle lysate was prepared from muscles of WT, *mdx* and p47^−/−^-*mdx* mice and subjected to immunoprecipitation using anti-Beclin, and the associated ATG14L, and Bcl-2 were determined using immunoblotting. Immunoblot for WT and *mdx* group are the same as shown in **Fig.1c**. **(b-c)** Immunoblot for JIP-1/JNK signaling proteins which participates in vesicle nucleation. Densitometry analysis of western blots represented by box plots **(d-e)** Protein expressions of autophagosome elongation marker, ATG5-12 was determined by western blot. Densitometry analysis of western blot represented by box plots. **(f)** Skeletal muscle tissue lysate was prepared from muscles of WT, *mdx* and p47^−/−^-*mdx* mice and subjected to immunoprecipitation using anti- SNAP29, and the associated SNAP29, STX17, ATG14L, and VAMP8 were determined using immunoblotting. Immunoblot for WT and *mdx* group are the same as shown in **Fig.2c. (g-h)** Immunoblots of lysosomal proteins, LAMP2 and Cathepsin-B. Densitometry analysis of western blot represented by box plots. GAPDH served as a loading control. (*n*=3 per group in all the figure panels). Statistical difference between groups were determined using ANOVA with Tukey’*s post hoc* test.

### Altered MTs acetylation affects autophagy

Emerging evidence shows that reversible acetylation of α-tubulin can regulate MT function, thus, facilitating fusion of autophagosomes to lysosomes^40, 41^. The acetylation status of microtubules is coordinated by deacetylases (HDAC6)^23^ and acetyltransferases (MEC-17) ^42^. Immunoblot analyses showed that the α-tubulin acetylation levels were significantly decreased in *mdx* muscles as compared to WT **(Fig. 4a-b)**. The deacetylase enzyme HDAC6 was increased in *mdx* muscle **(Fig. 4a-b)** whereas the acetyltransferase (MEC17) did not exhibit any change in *mdx* muscle **(Fig.S3a-b).** At the mRNA level, expressions of *hdac6, mec17* genes did not show any change in *mdx* as compared to WT **(Fig.S3c-d**). Autophagosome movement along MT tracks is dependent upon the microtubule motor proteins kinesin and dynein. Kinesin-1 (conventional kinesin or KIF5), is a tetramer of two kinesin heavy chains (KHC, KIF5B) and two kinesin light chains (KLC) ^43^ . We found that KIF5B was significantly increased in *mdx* muscles whereas KLC and dynein did not change in *mdx* as compared to WT (**Fig.4c-d).** KLCs play a dual role, they direct cargo binding and regulate motor activity. JIP-1 scaffolds with kinesin to link cargo to the kinesin-1 complex (KHC/KLC) ^44^. These observations raised the question of whether the JIP-Kinesin-1 complex is disrupted and thereby decreases binding to MTs and impairing autophagosome transport in *mdx* muscle. Surprisingly, JIP-1 was found to interact with both KLC and JNK in *mdx* mice similar to WT (**Fig.4e**). In addition, we did not find any restoration of acetylated α-tubulin levels in p47^-/-^/*mdx* mice **(Fig. 4f-g).** Overall, our data strongly suggests that microtubule acetylation plays an important role in autophagosome-lysosome fusion in *mdx* mice, which does not appear to be regulated by Nox2-ROS.

**Fig. 4.**
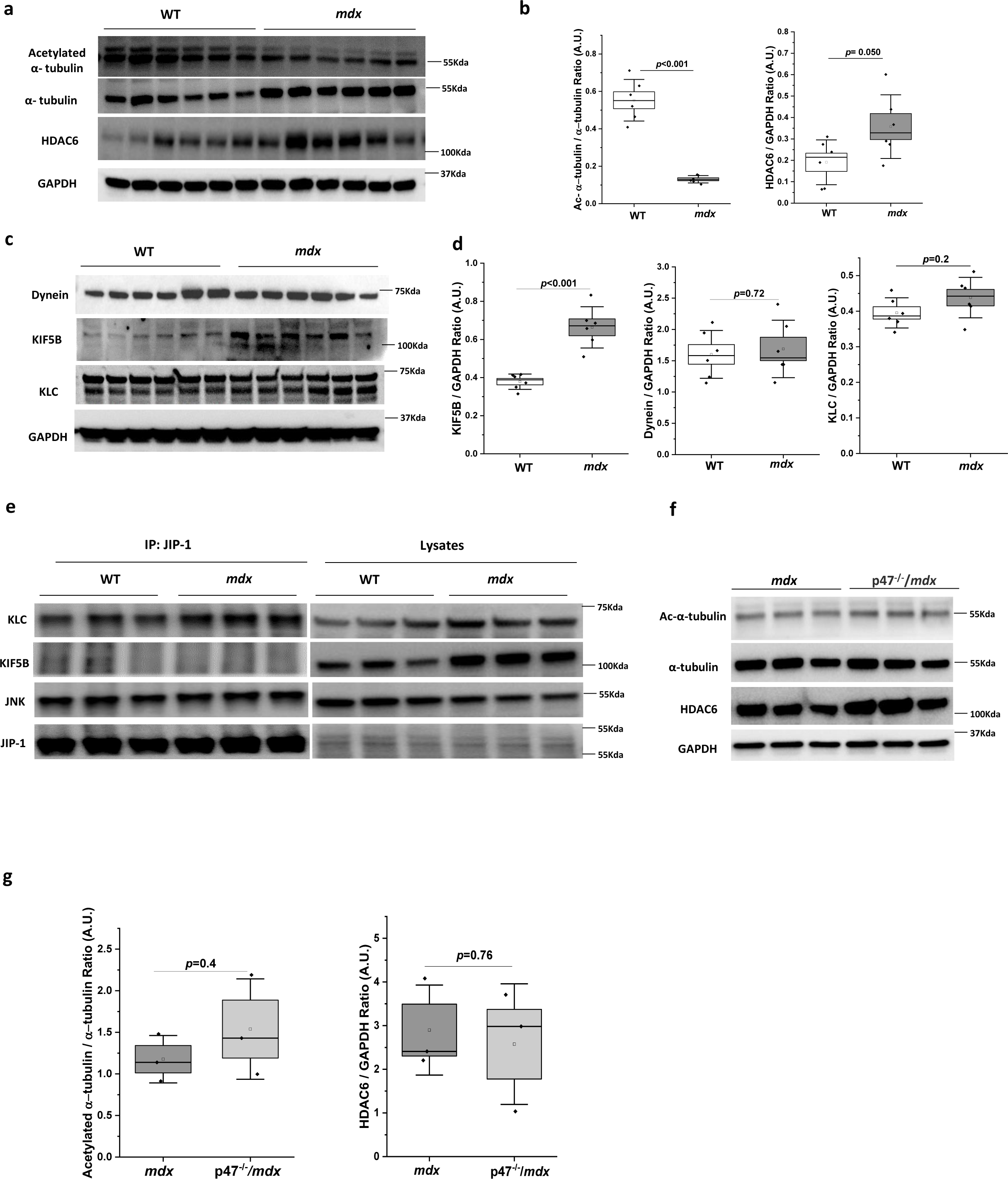
Altered tubulin-acetylation in *mdx* muscle. **(a-b)** Protein expressions of α-tubulin, acetylated-α-tubulin, HDAC6 were determined by western blot (n=6 per group). Densitometry analysis of immunoblot represented by box plots. **(c-d)** Protein expressions of MTs motor proteins, kinesin-1 complex (KIF5B, KLC) and dynein were determined by western blotting. WT and *mdx* mice (n=6 per group). Densitometry analysis of immunoblot represented by box plots **(e)** Skeletal muscle tissue lysate was prepared and subjected to immunoprecipitation using anti-JIP-1, and the associated KIF5B, KLC, and JNK were determined using immunoblotting (n=3 per group). **(f-g)** Protein expressions of α-tubulin, acetylated-α-tubulin, HDAC6 were determined by western blot in *mdx* and p47^-/-^/*mdx* mice (n=3 per group). GAPDH as a loading control. Statistical difference between two groups were determined using two-sample student-*t-*test.

### HDAC6 inhibition improves MT acetylation and promotes autophagosome-lysosome fusion in *mdx* muscle

Acetylated microtubules play an important role in vesicle trafficking and fusion. To assess the relationship of HDAC6 activity with microtubule alterations and impaired autophagic flux, HDAC6 was inhibited with its specific pharmacological inhibitor, Tubastatin A (TubA). TubA was intraperitoneally injected for 2 weeks in 3 week old *mdx* mice, which is just before the onset of disease progression. The dose given to the *mdx* mice was 70mg/kg per day which is equivalent to 8.4mg/kg per day for a human child based on FDA approved mouse to human-equivalence dose calculation guide^45^. TubA restored α-tubulin acetylation levels without altering the protein expression levels of either de-tyrosinated α-tubulin or HDAC6 **(Fig.5a-b)** nor kinesin, JIP-1, or JNK (**Fig.S4a-b**). We next assessed whether the increased acetylation of α-tubulin improved autophagic flux in *mdx* muscle. The autophagosome substrate p62 (SQSTM1) was significantly decreased in TubA treated *mdx* muscle while lipidation of microtubule-associated protein 1A/1B- light chain 3 (LC3II), which makes up the autophagosomal membrane showed only a small increase (**Fig. 5c-d**). To further examine the mechanism behind the efficient autophagic clearance in TubA treated *mdx*, we evaluated the autophagosome-lysosome fusion by detecting the binding of SNARE complex proteins. TubA treatment enhanced the interaction of SNAP29 with STX17 and VAMP8 **(Fig.5e).** Immunostaining of SNAP29 and VAMP8 showed puncta within the muscle and increased colocalization in TubA treated *mdx* muscle as compared to non-treated, indicative of enhanced autophagosome-lysosome fusion by HDAC6 inhibition **(Fig.5f-g).** In addition, the lysosomal protein LAMP2 and cleaved (active) lysosomal hydrolase cathepsin-B are elevated in TubA treated *mdx* muscle (**Fig. 5h-i**). Overall, our data suggest that the decreased acetylation of α-tubulin in *mdx* muscle inhibits autophagosome-lysosome fusion, which can be recovered upon HDAC6 inhibition.

**Fig. 5.**
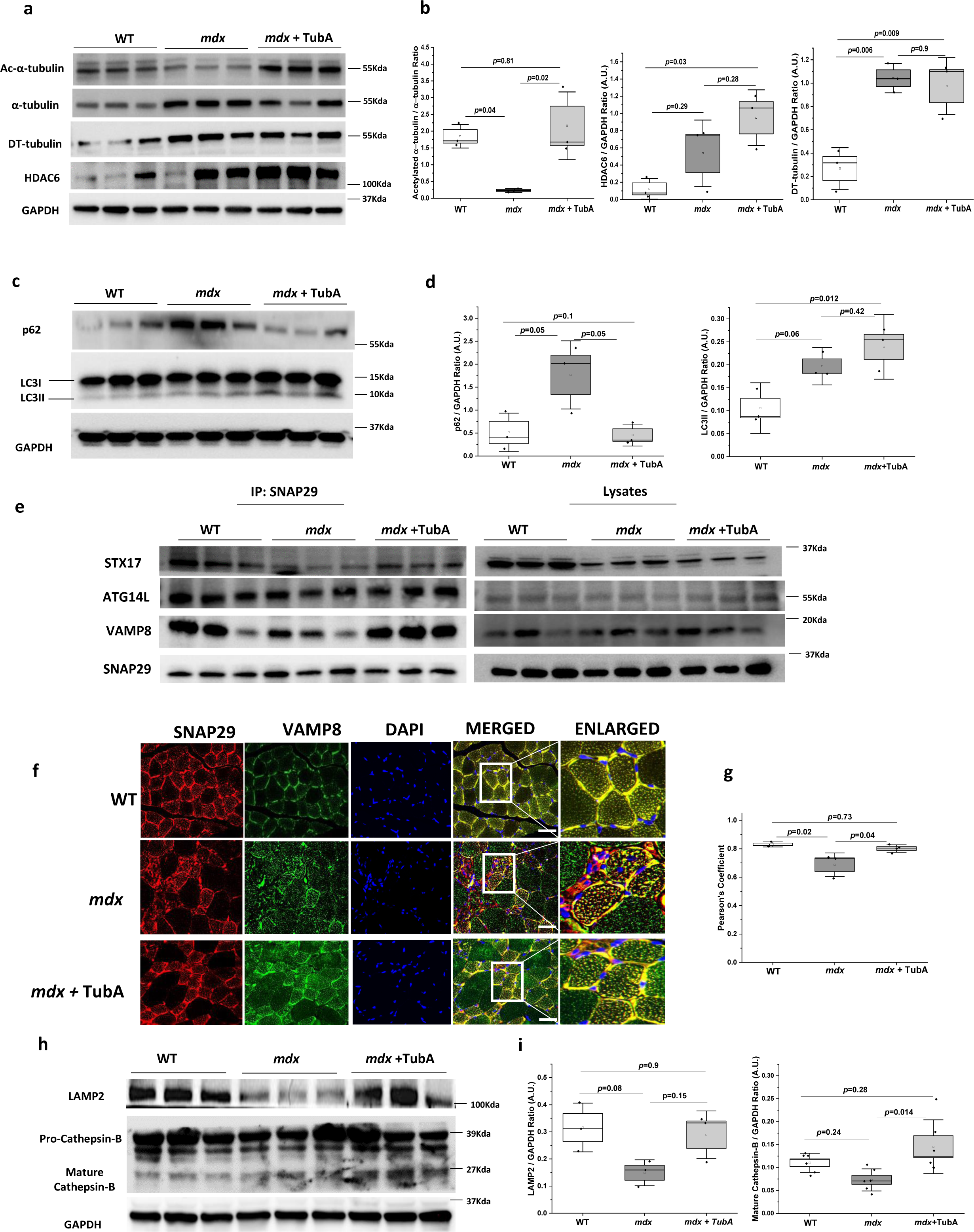
HDAC6 inhibition promotes tubulin acetylation and recovers autophagic flux in *mdx* mice. **(a-b)** Protein expressions of α-tubulin, acetylated-α-tubulin, detyrosinated α-tubulin, and HDAC6 were determined by western blot in WT, *mdx* and TubA treated *mdx* mice (n=3 per group). Densitometry analysis of immunoblot represented by box plots. **(c-d)** Immunoblot for autophagy related proteins- p62 and LC3 (n = 3 per group). Densitometry analysis of western blots represented by box plots. **(e)** Skeletal muscle lysate was prepared and subjected to immunoprecipitation using anti-SNAP29, and the associated, STX17, ATG14L, and VAMP8 were determined using immunoblotting (n=3 per group). **(f-g)** Representative immunofluorescence micrograph of anti-SNAP29 labeled with secondary antibody *Alexa Fluor 594 (Red) and* anti- VAMP8 labeled with secondary antibody *Alexa Fluor 488 (Green)*, counterstained with DAPI *(nuclear stain)* from section of gastrocnemius (GAS) muscle from WT (n=3), *mdx (n=3)* and TubA treated *mdx (n=5)*(*scale bar -50μm; Magnification-40X).*Quantitative values for colocalization analysis was represented as the Pearson’s Correlation Coefficient of SNAP29-VAMP8. **(h-i)** Protein expressions of lysosomal proteins, LAMP2 (n=3 per group) and Cathepsin-B ((n=6 per group) were determined by western blot in WT, *mdx,* and TubA treated *mdx* mice. Densitometry analysis of western blots represented by box plots. GAPDH served as a loading control. Statistical difference between groups were determined using ANOVA with Tukey’*s post hoc* test.

### HDAC6 inhibition recovers acetylation of PrxII and increases total PrxII in *mdx* mice

Prx I and Prx II are specific substrates of HDAC6, their acetylation status provides resistance to hyperoxidation. We have previously shown hyperoxidation and proteolytic degradation of PrxII in *mdx* skeletal muscle ^46^. Here, we immunoprecipated PrxII and probed for acetyl lysine and found decreased acetylated PrxII levels in *mdx* mice, which was recovered back to WT in TubA treated *mdx* mice (**Fig 6a**). This was further confirmed by immunoprecipitating acetyl lysine followed by probing for PrxII **(FigS5a).** Total PrxII levels were found to be significantly increased in TubA- *mdx* mice while oxidized PrxII (PrxSO2/3) showed no difference as compared to *mdx* **(Fig. 6b-c)**. Our findings suggests that TubA did not decrease PrxSO2/3 is not surprising, as sulfonation of PrxII is an irreversible oxidative modification ^47^ . We did find that the ratio of oxidized PrxII to total PrxII is significantly reduced in TubA-treated *mdx* (**Fig. 6b-c**), indicating that the increased acetylation of PrxII prevents its hyperoxidation. To our surprise, we found that stretch activated ROS production was not diminished in diaphragm muscle from TubA treated *mdx* compared to *mdx* (**Fig.6d**). Intriguingly, acetylated PrxII levels were restored back to WT levels in p47^-/-^/*mdx* mice (**Fig. S5b-c**). Total PrxII was increased, PrxSO2/3 decreased, and the ratio of oxidized to total PrxII decreased in p47^-/-^/*mdx* as compared to *mdx.* **(Fig. S5d-e).** These data suggest that either reducing oxidative stress (p47^-/-^/*mdx*) or increasing acetylation of PrxII (TubA) prevents its hyperoxidation and degradation in *mdx* skeletal muscle.

**Fig. 6.**
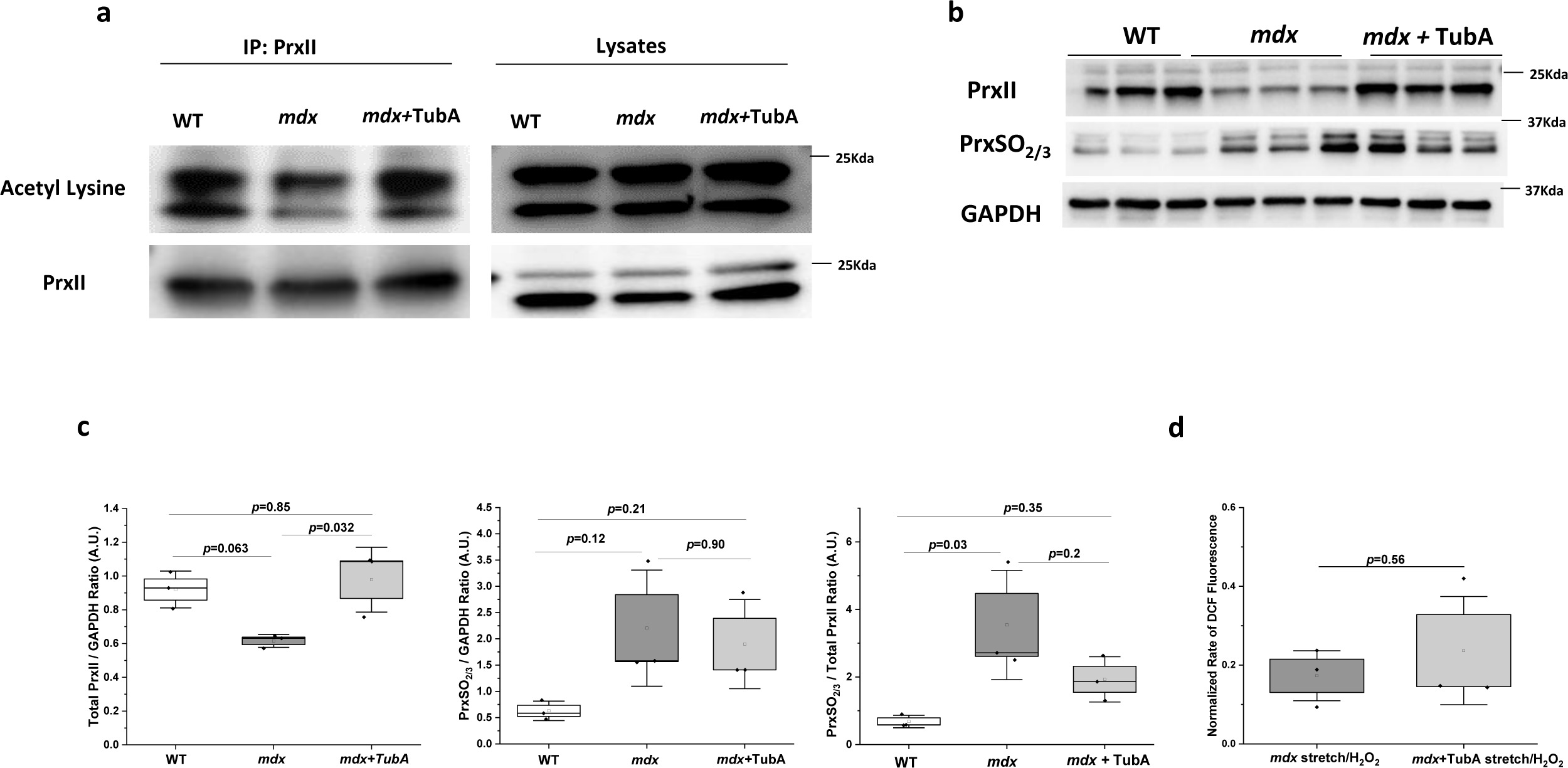
HDAC6 inhibition prevents deacetylation and hyperoxidation of PrxII in *mdx* mice. **(a)** Skeletal muscle lysate was prepared and subjected to immunoprecipitation using anti-PrxII, and the acetylation levels of PrxII was determined with the anti-acetyl lysine antibodies **(b-c)** Protein expressions of total PrxII and Prx-SO2/3 were determined by western blot in WT, *mdx,* TubA treated *mdx* mice (n=3 per group). Densitometry analysis of western blots and the ratio of PrxSO2/3 and total PrxII were represented by box plots. **(d)** Stretch-induced ROS measurements in diaphragm muscles of *mdx (n=2)* and TubA treated *mdx (n=3)* mice. GAPDH served as a loading control. Statistical difference between groups were determined using ANOVA with Tukey’*s post hoc* test. . Statistical difference between two groups were determined using two-sample student-*t-* test and Welch’s correction test was performed for non-equal variance between two groups in **panel d.**

### TubA treatment alleviates apoptosis and immune cell infiltration in *mdx* muscle

Impaired autophagy is associated with aggregation of oxidized/misfolded proteins and other cellular constituents, eventually leading to cell death. Immunoblot expression of cleaved/active caspase-3 was found to be significantly reduced in *mdx* mice following treatment with TubA **(Fig.7a-b).** The percentage of TUNEL positive nuclei (*green, marked by white arrows*) were significantly elevated in *mdx* gastrocnemius (GAS) muscle, which were significantly decreased upon treatment with TubA (RITubA =92.14%, **Fig. 7c-d**). Infiltration of macrophages and other immune cells is an important and pathogenic feature of dystrophin-deficient muscle, even at asymptomatic stage of disease progression. We have stained GAS muscle cross-sections with anti- CD68 antibody to identify CD68+ macrophages. We found a significant decrease in CD68^+^ immune cells *(green, marked by white arrows)* in TubA treated *mdx* skeletal muscle (RI TubA=63.10%, **Fig. 7e-f**), indicating decreased infiltration of macrophages in the endomysium of skeletal muscles.

**Fig. 7.**
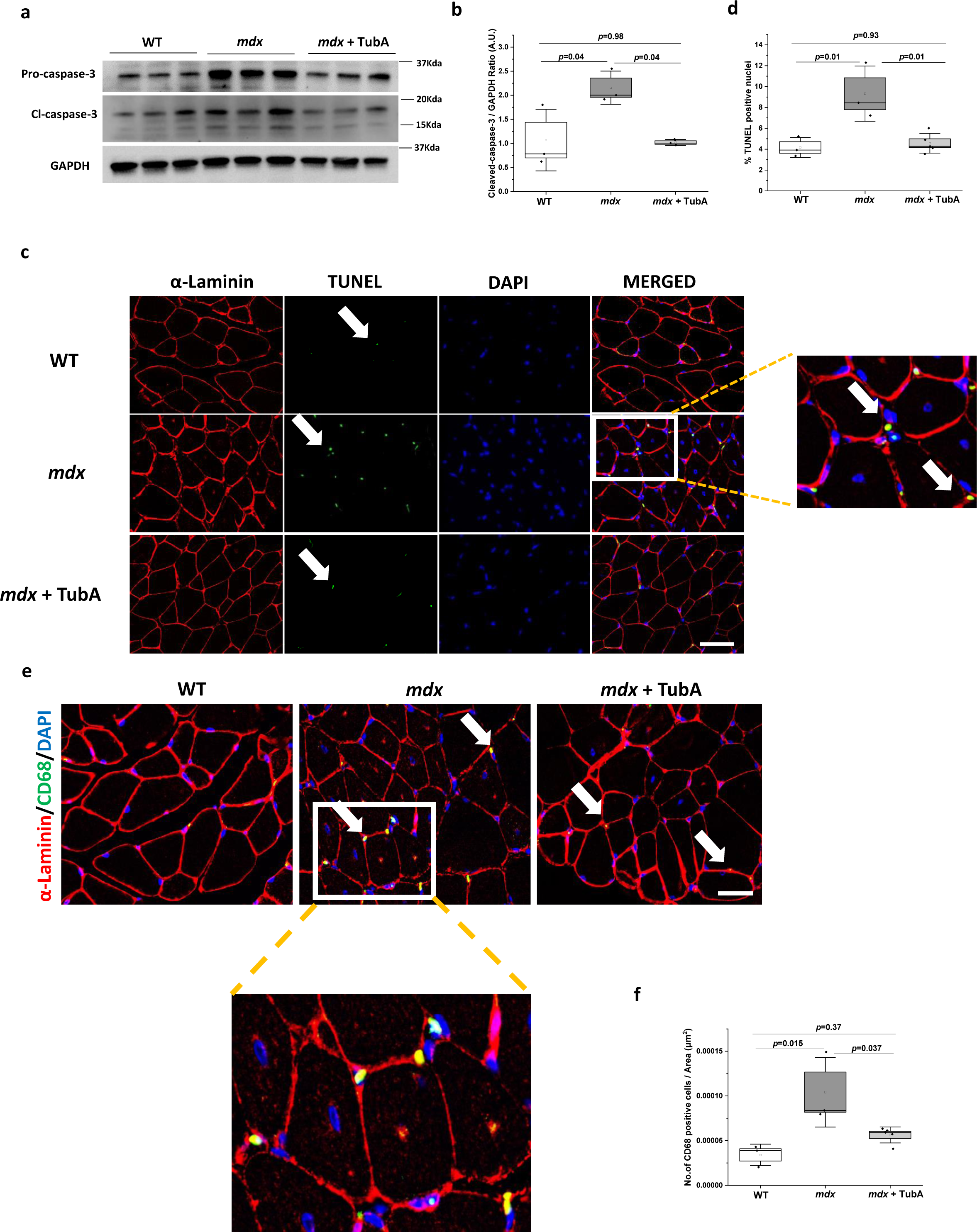
TubA treatment alleviates apoptosis and immune cells infiltration in *mdx* muscle. **(a-b)** Protein expressions of pro-caspase-3 and cleaved caspase-3 were determined by western blot from TA muscles extract of WT, *mdx*, and TubA treated *mdx* (n = 3 per group). Densitometry analysis of immunoblots represented by graph. GAPDH as a loading control **(c-d)** Paraffin- embedded gastrocnemius (GAS) muscle sections (4µm) of WT (n=3), *mdx (n=3)* and *mdx*+TubA (n=5) mice processed for the detection of TUNEL positive nuclei *(green).* Sections were stained with α-laminin *(red)* to define the sarcolemma. Nuclei were counterstained with DAPI *(blue).* White boxed region shows the enlarged image from the *mdx* muscle section in the panel on right indicating TUNEL positive nuclei (*white arrow)*.Numbers of TUNEL positive nuclei counted by Image J software. **(e-f)** Macrophage infiltration analyzed by Immunofluorescence staining of CD68 (*green*), α-laminin (*red*), and nuclei (*blue*) in GAS section isolated from WT (n=3), mdx (n=3), *mdx*+TubA (n=5) mice. White boxed region shows the enlarged image from the *mdx* muscle section indicating infiltration of CD68+ cells in the skeletal muscles *(white arrow)*.Quantification of CD68 positive immune cells. *Scale bar (50µm), Magnification-40X*. Statistical difference between groups were determined using ANOVA with Tukey’*s post hoc* test.

### Amelioration of dystropathology and improvement in muscle functional assessment following TubA treatment

Since pharmacological inhibition of HDAC6 improved MT acetylation, restored autophagic flux by promoting autophagosome-lysosome fusion, and decreased inflammation and apoptosis we next investigated whether TubA reduced muscle damage and improved muscle performance. Treatment of *mdx* mice with TubA prevented the increase in serum creatine phosphokinase (CPK activity (RITubA=133.81%, **Fig. 8a**), a widely used clinical marker of muscle damage. The etiology of DMD can be explained by loss of membrane integrity due to dystrophin deficiency which results in degeneration of myofibers. To determine whether TubA affects sarcolemmal integrity, we intraperitoneally injected mice with Evans blue dye (EBD), which penetrates non-specifically into any cell with disrupted/leaky membranes. At 24 h after injection, EBD accumulated abundantly in *mdx* muscles as compared to WT muscles **(Fig. 8b),** confirming membrane permeability due to the loss of dystrophin. Notable, TubA treatment significantly blunted EBD uptake into the DIA muscle of *mdx* mice. This observation was further confirmed by fluorescence staining of cross-sections of DIA muscles, the number of EBD positive fibers were reduced in *mdx* mice treated with TubA (RI TubA=85.81%, **Fig.8c**).

**Fig. 8.**
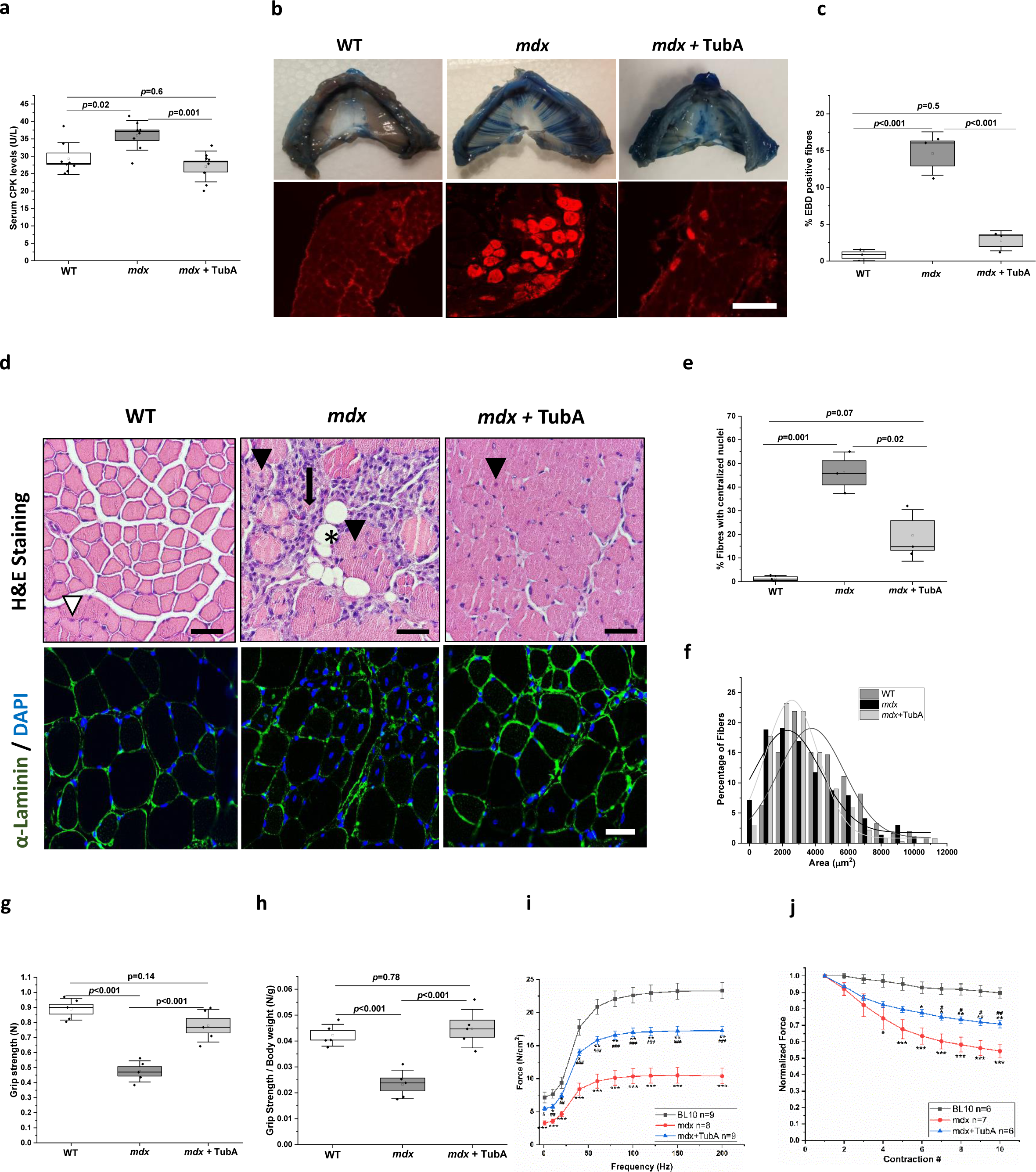
HDAC6 inhibition ameliorates muscle pathophysiology and improves muscle function. **(a)** Serum creatine phosphokinase quantitated by ELISA (n = 8 per group) **(b-c)** Loss of sarcolemmal integrity was evaluated by the *i.p*. Injection of Evans Blue Dye (EBD) in mice. Dye injected 24hr prior to the completion of two weeks TubA treatment. EBD staining of diaphragm (DIA) muscles and its cross section (4μm) showing EBD positive fibers. Quantification of EBD positive fibers in DIA muscle fibers (n=3 per group). *Scale bar (90µm), Magnification-20X.* **(d)** H&E-stained muscle (TA). White arrow head= peripheral nuclei; black arrow head=central nuclei; black arrow=necrotic myofibers (infiltration of immune cells); black asterisk=fat depositions within myofibers. (n = 3 mice per group), *Scale bar: 50 μm, Magnification-40X*. Immunofluorescence staining of α-laminin (*green)* and DAPI (*blue nuclei staining*) in TA cross-section muscle of WT, *mdx*, TubA treated *mdx*. (n=3 per group) (e) Percentage fibers with centralized nuclei was quantified by Image J. *Scale bar: 50 μm, Magnification-40X* **(f)** Myofibers CSA calculated from minimum feret’s diameter (n=3 per group). **(g-h)** Grip strength, absolute and normalized to body weight (n = 5 per group). **(i)** Force–frequency relationship in DIA muscle strips from WT (n = 9), *mdx* (n = 8), and *mdx*+TubA (n = 9). **(j).** Eccentric contraction induced force drop (normalized to the first contraction) in EDL muscles isolated from WT (n=6), *mdx (n=7)*, TubA treated *mdx (n=6)* mice. Statistical difference between groups were determined using ANOVA with Tukey’*s post hoc* test.*p<0.05 vs BL10, **P<0.01 vs BL10, ***p<0.001 vs BL10, *P<0.05 vs mdx, **p<0.01 vs mdx for panel i and j.

We next assessed the histopathological features of TA muscles from WT, *mdx* and TubA treated *mdx* mice by H& E staining. We observed severe necrosis (*marked by black arrow*), regenerating fibers with central nuclei *(marked by black arrow head)*, and fat deposition in the interstitial space *(marked by black asterisk)* of dystrophic muscle as compared to WT muscles. Notably, overall necrotic myofibers, central nuclei, and fat depositions were reduced in TubA treated *mdx* **(Fig.8d).** We further quantified histological sections of TA muscles by immunostaining with α-laminin (*green*) and DAPI (*blue)* to analyze the cross-sectional area (CSA) and centronucleated myofibers in treated and untreated *mdx* mice **(Fig.8d)**. A drastic increase in the percentage of fibers with centralized nuclei was observed in *mdx* TA muscle as compared to WT (46% versus 1.2%), which was significantly reduced in TubA-treated *mdx* muscle (RITubA= 52.2%, **Fig. 8e)**. TA muscle from *mdx* mice showed decreased cross-sectional area (CSA) as compared to WT, which was partially prevented with TubA treatment as exhibited by the distribution of Minimal Feret’s diameter **(Fig 8f).** DMD patients suffer from progressive muscle weakness, eventually leading to immobility and respiratory failure. To examine whether TubA improves muscle function and strength, *in-vivo* muscle functional performance was assessed by grip strength. Grip strength was conducted after completion of 2weeks of TubA treatment in *mdx* mice. Skeletal muscle strength was significantly improved in TubA treated *mdx* mice as compared to non-treated *mdx* mice (RITubA**=**113.29%, **Fig. 8g-h).** Finally, to determine the effects on contractile function, we compared the force-generating capacities of untreated and TubA-treated *mdx* diaphragm muscle strips electrically stimulated *ex vivo.* TubA-treated *mdx* mice demonstrated significantly greater diaphragmatic force production over a broad range of stimulation frequencies (0 to 200 Hz) **(Fig. 8i, Table 1).** In addition, TubA treated *mdx* EDL muscle was significantly protected from eccentric contraction induced force loss compared to *mdx* muscle **(Fig. 8j, Table 1).**

**Table 1.** Recovery Index for Tubastatin A treatment of mdx.

| Outcome Measures | Recovery Score by TubA (70mg /kg b.w. for 2 weeks) in *mdx* mice |
| --- | --- |
| Serum Creatine phosphokinase (CPK) | 133.8% |
| Immune Infiltration (CD68<sup>+</sup> cells) | 63.10% |
| Centralized nuclei | 52.2% |
| Apoptotic cell death | 92.14% |
| EDL ECC force production at 10<sup>th</sup> contraction | 46.68% |
| Diaphragm Isometric force production (200Hz) | 53.24% |
| Grip Strength normalized with Body weight | 113.29% |

## Discussion

Previous studies have shown that defective autophagy and disorganized MT network play an important role in disease progression and contributes to DMD pathogenesis before the manifestation of severe phenotypes ^11, 48–51^. Therefore, reverting autophagic dysfunction could be an efficient approach to slow the onset of disease progression and improve the muscle function in DMD patients. Our group was the first to identify that Nox2 mediates MT alterations and autophagic dysfunction in dystrophic muscle ^7, 14^. MTs regulate autophagy through their scaffolding and transport functions. Acetylated MTs induce autophagy by facilitating intracellular trafficking and fusion of autophagosomes with lysosomes^18, 19, 52^. Despite the number of investigations, no consensus about the role of tubulin post translational modifications in regulating autophagic flux in DMD have been reached. We are the first to report that autophagy is differentially controlled by redox and acetylation modifications in *mdx* mice (**Fig. 9**). We provide evidence that genetic ablation of Nox-2 in *mdx* mice promotes autophagosome maturation without facilitating autophagosome-lysosome fusion. Further, our study reveals alterations in MT acetylation in dystrophin-deficient muscle, which is not restored back in Nox-2 ablated *mdx* mice. Pharmacologically targeting HDAC6 restored MT acetylation, rescued autophagy by accelerating fusion and content degradation, restricted muscle damage and improved muscle function in *mdx* mice.

**Fig. 9.**
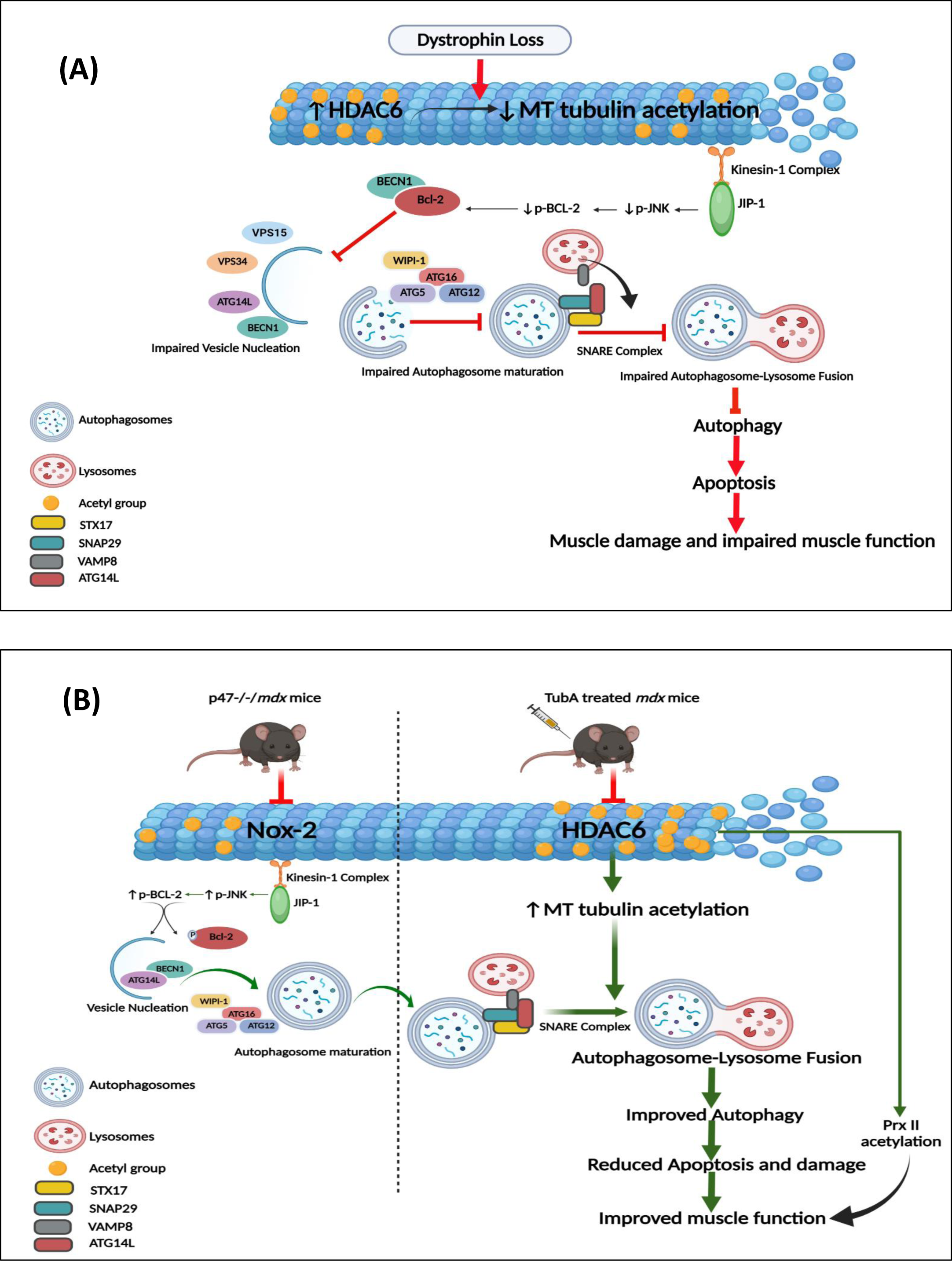
Model of Impaired autophagy in mdx skeletal muscle are differentially regulated by redox and acetylation modifications. **(A)** Loss of dystrophin leads to increased HDAC6 expression and subsequent deacetylation of α-tubulin. Decreased α-tubulin acetylation inhibits binding of the kinesin-/JIP-1 complex and subsequent transport of autophagosomes to lysosomes. Decreased activation of JIP-1 results in decreased phosphorylation of JNK and Bcl-2, preventing the dissociation of BECN1/Bcl-2 and sequestering BECN1 away from the PI3classIII complex (ATG14L-Vps34-Vps15). The net result is impairment of phagophore nucleation. Decreased WIPI-1 inhibits localization of ATG5-12/16L1 complex to the phagophore, leading to impaired autophagosome maturation in mdx mice. SNARE associated proteins play a major role in membrane-mediated events of autophagosome-lysosome fusion, another crucial step in autophagy process. In mdx mice, reduced interaction of SNAP29 with STX17 and VAMP8 leads to the inhibition of autophagosome-lysosome fusion. **(B)** Genetic inhibition of Nox-2 activity in mdx mice activates JIP-1, resulting in phosphorylation of JNK and Bcl-2, dissociation of BECN1 from Bcl-2 and induction of phagophore nucleation. In addition, Nox-2 inhibition promotes ATG5-12 complex formation and therefore autophagosome maturation. However, autophagosome-lysosome fusion is not improved upon inhibition of Nox-2 activity, likely due to no change in α-tubulin acetylation. Pharmacological inhibition of HDAC6 promotes α-tubulin acetylation and improves SNARE complex formation, thereby facilitating fusion of autophagosomes with lysosomes, improves autophagy, decreases dystropathology and improves skeletal muscle function. Increased acetylation of the antioxidant enzyme Prx II likely contributes to improved muscle function in mdx skeletal muscle. **Figure created with <u>BioRender.com</u>**

Autophagosome biogenesis/maturation includes phagophore nucleation, expansion, and closure of phagophore membranes, generating the complete autophagosome^36^. Initiation of autophagy requires the PI3KClassIII complex consisting of VPS34, VPS15, Beclin1/BECN1 and ATG14L. Emerging evidence suggests impaired autolysosome clearance in VPS15 deficient muscle and the muscle phenotype evocative of lysosomal myopathies ^53^. A study by Nascimbeni et al., have shown that alterations in VPS15, VPS34 and BECN1 lead to impaired endosome and lysosomal maturation in Danon Disease and Glycogen Storage Disease (GSDII) ^54^. In the current study, we found that the PI3KclassIII complex was disrupted in *mdx* muscle, with a significant decrease in ATG14L and VPS34, whereas VPS15 was found to be increased. Intriguingly, we also found reduced interaction of Beclin-ATG14L complex whereas Beclin-Bcl-2 complex formation, which is known to inhibit autophagic induction, is increased in *mdx* muscle. This indicates disruption in the formation of vesicle nucleation. During starvation, tubulin acetylation signals kinesin recruitment to the MTs, with subsequent JNK activation and phosphorylation of Bcl-2, which allows the release of Beclin from Beclin–Bcl-2 complexes to initiate autophagosome formation.^55^. We found decreased JNK activation, likely due to reduced levels of the scaffolding protein JIP- 1, in *mdx* muscles and reduced phosphorylation of Bcl-2 (S70), thus inhibiting Beclin-ATG14L association for vesicle nucleation. In addition, WIPI-1, which recruits lipid phosphatidylinositol- 3 phosphate (PI3P) and mediates recruitment of the ATG5-12/16L1 complex and the LC3 lipidation, was found to be reduced in skeletal muscle from *mdx* mice. Taken together, our data suggest that defects in autophagosome maturation is mediated by inhibition of the Kinesin-JIP- JNK pathway in dystrophic mice.

SNARE associated proteins play a major role in membrane-mediated events of autophagosome- lysosome fusion, another crucial step in autophagy process. Recent studies have highlighted the role of STX17-SNAP29-VAMP8 Q-SNARE tertiary complex in promoting autophagosome- lysosome fusion. ^19, 56^ We revealed that the blockage of autophagosome-lysosome fusion in *mdx* mice was not due to altered expressions of SNARE associated proteins but due to the inhibition of the interaction of STX17 and VAMP8 with SNAP29, which culminates in accumulation of autophagosomes ^7^

Previous work from our lab has shown that genetic ablation of Nox2-ROS improved autophagy in *mdx* mice. Nevertheless, the detailed mechanisms by which autophagy was rescued in the p47^-/-^ /*mdx* mouse was not evaluated, which is required to improve the therapeutic outcomes for DMD patients. Our current data showed that p47^-/-^/ *mdx* mice prevented the defects of autophagosome maturation by promoting vesicle nucleation and elongation of autophagosomes. However, our data revealed that genetic inhibition of Nox-2 was not able to promote the interaction between STX17- SNAP29 and VAMP8-SNAP29 and thus cannot promote autophagosome-lysosome fusion and content degradation in *mdx* mice. These finding may explain why we only observed partial rescue of dystrophic pathology in the p47^-/-^/*mdx* mouse^7^ .

Based on the above findings, we speculate that autolysosome formation in *mdx* muscle is redox independent, leading us to explore the mechanisms that underlie the failure of delivery of autophagosomes and its fusion with lysosomes. Protein acetylation controls autolysosome formation and improves autophagic flux^19, 39, 57^. Emerging evidences suggests that HDAC2 promotes autophagy by causing deacetylation of STX17, which increases its binding with SNAP29 and promotes the Q-SNARE complex formation ^39^. Interestingly, our findings revealed the deacetylation of STX17 and increased HDAC2 in *mdx* mice, which suggests that reduced interaction of STX17-SNAP29-VAMP complex is independent of the STX17 acetylation and HDAC2.

Previous studies from our lab and others have reported dystrophin deficiency results in disorganized MT lattice network and elevated MT tubulin modifications (detyrosinated α-tubulin) in *mdx* muscle ^14, 58^ . Post translational modifications of MTs, in particular acetylation, have been shown to facilitate autophagosome formation and serve to direct mature autophagosomes for fusion and degradation^41, 59, 60^. In the present study, we find decreased acetylated α- tubulin in *mdx* mice, with increased protein expressions of the deacetylase enzyme, HDAC6. Accumulating evidence suggests that JIP-1 mediates and regulates the binding of cargo to the Kinesin-1 complex via KLC on the microtubule track. Activated JNK regulates the binding of cargo to KLC by mediating the release of cargo once they have reached the lysosomes for fusion. Ittner et al. have shown that impaired binding of JIP-1 to the kinesin-1 complex disrupted the anterograde axonal transport of cargo along MTs in Alzheimer’s disease^61^. Surprisingly, in dystrophic muscle, JIP-1 interacts with KLC and JNK, forming a KLC-JIP-JNK complex. We therefore speculate that decreased MT acetylation in *mdx* muscle can prevent the binding of KLC-JIP-JNK on the MT track, leading to inhibition of autophagosome transport and its fusion with lysosomes.

HDAC6 inhibition exerted beneficial effects by improving autophagic dysfunction through MT acetylation in several pathophysiological conditions such as, Huntington’s disease (HD)^27^spinal cord injury ^19^, osteoarthritis ^33^, and doxorubicin-induced cardiomyopathy^30^. HDAC6 has unique structure and properties, exerting both enzymatic and non-enzymatic effects on cellular functions. In addition to two catalytic domains which deacetylates cytoplasmic non-histone proteins, it also possesses a non-enzymatic zinc-finger ubiquitin-binding domain (UBD) at its C-terminus^62^. Through its UBD, HDAC6 interacts with ubiquitinated protein aggregates to promote loading and transport along the microtubules for proper degradation. Given this dual role of HDAC6 in regulating the response to cellular stress, pharmacological inhibition of its enzymatic activity, as opposed to knock-down of protein content, has been shown to be a novel and promising therapeutic strategy for several diseases^63^. We selected TubA, which stands out as a potent and highly specific HDAC6 inhibitor with an IC50 of 15 nM and has a strict selectivity for HDAC6 over all other HDACs (over 1000-fold) except for HDAC8 (57 folds) ^64^. Unlike other HDAC inhibitors, TubA exhibits no toxicities such as fatigue, nausea, or thrombocytopenia, thus making HDAC6 a most suitable and promising therapeutic target ^64^. We assessed TubA dosing in *mdx* cohorts at 30mg/kg every other day for 4weeks and did not observe any muscle functional recovery, although serum CPK levels were reduced. **(Fig.S6a-b**).We further assessed TubA dosing in *mdx* cohorts at 70mg/kg every day for 2 weeks. Interestingly, TubA treatment at this dose showed improvement in muscle contractile properties and strength, and reduced serum CPK levels in *mdx mice*. Our data shows that HDAC6 inhibition specifically promoted tubulin acetylation without affecting DT- tubulin levels in *mdx* mice. However, HDAC6 inhibition was not able to activate JIP/JNK signaling as assessed by phospho-JNK and thus, autophagosome maturation was not improved in TubA treated *mdx*. HDAC6 inhibition was recently reported to restore neuronal autophagy flux by increased LC3II and decreased p62 aggregates in neurodegenerative disorders^27, 65^. TubA treatment promoted the autophagosome-lysosome fusion by SNARE machinery in SCI^19^. Consistent with these findings, we found increased LC3II and reduced p62 aggregates, which indicates efficient autophagic clearance in TubA treated *mdx.* TubA treated *mdx* mice showed increased interaction of STX17 and VAMP8 with SNAP29, thus facilitating the fusion and formation of autolysosomes, as well as enhanced lysosomal function. Altogether, our data support a role for HDAC6 regulating autophagic flux by decreasing MT acetylation and inhibiting protein aggregate trafficking and fusion of autophagosome-lysosome in skeletal muscle from *mdx* mice, which is reversed upon HDAC6 inhibition with TubA.

HDAC6 may have a role in regulating redox balance within the cell, as HDAC6 has been shown to deacetylate redox regulatory proteins, PrxI and PrxII^25^; and once deacetylated PrxI and PrxII lose their antioxidant capacity. HDAC6 inhibition recovered acetylation of Prx and provided protection against oxidative stress in Aβ-induced Alzheimer’s disease (AD) ^24^, 6-OHDA induced Parkinson’s disease (PD) model ^29^, and myocardial ischemia/reperfusion (MI/R) ^32^, suggesting a new mechanism regulating redox balance. In the current study, our results showed that acetylation levels of PrxII were decreased in *mdx* muscle, and TubA treatment recovered both acetylation levels of PrxII and total PrxII in *mdx* mice. Our previous findings show that PrxII overexpression protects *mdx* mice from eccentric contraction induced force loss^46^. However, we did not observe any change in either oxidized PrxII (PrxSO2/3) or stretch dependent ROS production in *mdx* mice treated with TubA. Intriguingly, p47^-/-^/*mdx* mice showed reduced levels of oxidized PrxII (PrxSO2/3), increased PrxII acetylation, but did not increase MT acetylation. Thus, there appears to be cross-talk between HDAC6 and Nox2 in dystrophic skeletal muscle that we have yet to understand, and doing so will be essential for developing effective therapeutics for maintaining muscle homeostasis in DMD patients.

Our data reveals that TubA has the potential to act on the downstream pathogenic events of dystropathology. TubA treatment attenuated apoptosis, decreased levels of CD68+ macrophages infiltrated into the muscle, maintained sarcolemmal integrity, and remarkably reduced serum CK levels in *mdx* mice. Quantitative analysis of Hematoxylin/eosin staining showed significant reduction in the centralized nuclei and increased CSA in TA *mdx* muscle following treatment with TubA. In DMD patients, even at very young ages, weakness in hand strength is observed as a generalized effect of dystrophin deficiency. Loss of grip strength was observed in *mdx* mice while TubA treated *mdx* mice showed significant improvement in muscle strength. Diaphragm is the most severely affected skeletal muscle in dystrophic mice, and diaphragm dysfunction is a major cause of respiratory failure in DMD patients. TubA-treated *mdx* mice showed significantly greater diaphragm force production and decreased ECC induced force loss. These finding indicate that HDAC6 inhibition not only restores autophagy flux by regulating MT acetylation but it extended its efficacy to alleviate many symptoms due to loss of dystrophin in DMD In conclusion, our study suggests that defects in autophagy are regulated by both redox and acetylation modifications in *mdx* mice. Evidence provided in this study confirms the role of HDAC6 in deacetylating MTs, which led to the disruption of autophagosome-lysosome fusion in dystrophin deficient mice. HDAC6 inhibition holds the potential to counteract the alteration of the MTs and restore impaired autophagic flux in dystrophic skeletal muscle **(Fig.9).** This pre-clinical study suggests that targeting specific histone deacetylase, HDAC6 served as an effective tool in preventing disease progression and improving muscle strength and function **(Table 1)**, supporting the translational potential of TubA into clinical research for DMD patients. In view of our data, combinatorial therapeutic approaches are currently underway in our laboratory to assess the efficacy of TubA and Nox-2 pharmacological inhibitor in ameliorating pathological changes in DMD.

## Material and Methods

### Antibodies and Reagents

Tubastatin A and PEG300 were purchased from Selleck Chem, DCFH-DA (6-carboxy-2′,7′- dichlorodihydrofluorescein diacetate) was from Invitrogen. DMSO and Evans Blue Dye were purchased from Sigma Aldrich. Tween-80 from Fischer Scientific. Protein A Magnetic Beads, Mini-Protean TGX stain free gels, and Immun-Blot® PVDF Membrane, Clarity western ECL substrate were purchased from Bio-Rad. Anti-beclin, anti-p-bcl-2, anti-vps34, anti-vps15, anti-JNK, anti-p-JNK, anti-atg5-12, anti-atg7, anti-atg4b, anti-α-tubulin, anti-HDAC6, anti-p62, anti-caspase-3, and anti-LC3 antibodies were from Cell Signaling Technology. Anti-GAPDH (glyceraldehyde-3-phosphate dehydrogenase) and anti-dynein were purchased from EMD Millipore. Anti-CD68, anti-LAMP2, anti-Cathepsin-B, anti-SNAP29 were purchased from Santa Cruz Biotechnologies. Anti-WIPI1, anti-snap29, anti-stx17, anti-vamp8, anti-KIF5B, anti-KLC, anti-detyrosinated, anti-Prx-SO3, and anti-α-laminin were from Abcam. Anti-acetylated tubulin, anti-bcl-2, anti-atg14, anti-MAPKK, anti-JIP-1, anti-prxII were from Millipore Sigma. Secondary antibodies for immunofluorescence (Alexa Fluor® 594 donkey anti-rabbit, Alexa Fluor® 488- donkey anti-rabbit, Alexa Fluor®594-goat anti-mouse), and ProLong™ Gold Antifade Mountant with DAPI were purchased from Invitrogen. Microscopic slides was purchased from Denville Scientific Inc. Detailed information about antibodies and dilution can be found in **Supplementary Table 1 and 2.**

### Mice

C57BL/10ScSnJ (WT) and C57Bl/10ScSn-Dmd*mdx*/J (*mdx*) were purchased from Jackson Laboratories (Bar Harbor, ME) and bred following their breeding strategy. Mice with a genetic deletion of the Nox-2 scaffolding subunit p47^phox^ (p47^−/−^) were obtained from The Jackson Laboratory (B6 (Cg)-Ncf1m1J/J) and bred onto the *mdx* background as previously described ^7^. All animals used for experiments were males between 3-6 weeks of age. All animal studies were conducted in accordance with ARRIVE (Animal Research: Reporting of In Vivo Experiments). Mice were housed in a specific pathogen free (SPF) facility on a 12hr light/dark cycle. All mice were monitored daily and drug tolerability evaluated on the basis of body weight and clinical signs. After treatment period, mice were sacrificed by deep anesthesia followed by cervical dislocation as approved by the Institutional Animal Care and Use Committee (IACUC) of Baylor College of Medicine. Skeletal muscles were quickly excised, and either placed in Krebs-Ringer buffer for tissue experiments, snap frozen in liquid nitrogen for biochemical and molecular experiments, or in 10% formalin for histological experiments. We employed basic methods to analyze the preclinical experiments (TREAT-NMD) ^66^

### HDAC6 Inhibitor treatment

For pharmacological inhibition of HDAC6, HDAC6-Specific inhibitor TubastatinA (TubA) (Selleck-Chem, USA) was dissolved in 4% dimethyl sulfoxide (DMSO), 30% PEG300 and 66% MilliQ with a final concentration of 2.5 mg/mL, and was freshly prepared each day. TubA was intraperitoneally *(i.p.)* injected into 3 week-old *mdx* mice (onset of disease progression) at the dosage of 70 mg/kg b.w. once a day for 2 weeks.

### Fore-limb Grip strength analysis

Fore-limb grip strength was performed to assess the effects of TubA on recovery of muscle strength of *mdx* mice. The grip strength meter (Columbus Instruments, OH) was positioned horizontally and mice were held by the tail. Mice were allowed to grasp the smooth metal rod with their forelimbs only and then were pulled backward by the tail. The force applied to the metal rod at the moment the grasp was released and recorded as the peak tension. The test was repeated 5 consecutive times within the same session and the average value from the 5 trials was recorded as the grip strength for that animal. In order to avoid modifications induced by the body weight of the mice in the latency of staying on the grip strength meter, the data have been normalized with body weight of mice. The grip strength test was conducted by the same investigator in order to avoid variability in performing experiment, and the investigator was blinded to the treatment group.

### Serum Collection

Blood was collected from the hepatic portal vein of mice immediately following sacrifice and left at room temperature for 30 min to achieve coagulation. Serum was then separated from other blood fractions by centrifugation at 1,000 g for 10 min and stored at −80 °C for further use. CK activity in the serum was measured using Creatine Kinase Activity Assay Kit (My BioSource) following the manufacturer’s instruction.

### Ex-Vivo EDL Eccentric force measurements

Muscle contractile force measurement was conducted by using an *ex-vivo* muscle test system. In brief, the mice were anesthetized with isoflurane, the hind limb was excised and immediately placed in a bicarbonate-buffered solution (120 mM NaCl, 4 mM KCl, 1 mM MgSO4, 1 mM NaH2PO4, 25 mM NaHCO3, and 2 mM CaCl2, 10mM glucose) equilibrated with 95% O2-5% CO2 (pH7.4) for dissection. The proximal and distal tendons were tied with braided silk suture thread (4-0, Fine Science Tools) and mounted in a muscle bath containing bicarbonate-buffered solution continuously bubble with 95%O2, 5% CO2 between a fixed hook and a dual-mode lever system (305C-LR-FP, Aurora Scientific Inc., Aurora, ON, Canada) and allowed to equilibrate to 30°C for 10 minutes. The stimulation protocol consisted of supramaximal electrical current delivered through platinum electrodes using a biphasic high-power stimulator (701C; Aurora Scientific). Optimum length (Lo) was determined with a series of twitch stimulations measured using a hand-held electronic caliper, after which the muscle underwent 10 eccentric contraction with 3 minutes rest between each contraction so as to not elicit muscle fatigue. Each eccentric contraction consisted of a 200 ms isometric tetanus (150Hz) followed by stretch from 100% to 110% of Lo at 0.5Lo/s and then shortened to Lo passively. Following the 10 eccentric contractions the muscle was removed from the organ bath, trimmed of connective tissue, blotted dry, and weighed. Data were analyzed using the dynamic muscle control and analysis software (Aurora Scientific Inc.).

### Ex-Vivo Diaphragm Isometric force measurements

Diaphragm (DIA) muscle was dissected from mice and one end tied to a fixed hook and the other to a force transducer (F30, Harvard Apparatus) using silk suture (4-0) in bicarbonate-buffered solution continuously gassed with 95% O2–5% CO2 at 30°C. Contractile properties were assessed by passing a current between two platinum electrodes at supramaximal voltage (PanLab LE 12406, Harvard Apparatus) with pulse and train durations of 0.5 and 250 ms, respectively. Muscle length was adjusted to elicit maximum twitch force (optimal length, Lo) and the muscle was allowed a 10-min equilibration period. To define the force-frequency characteristics, force was measured at stimulation frequencies of 1, 5, 10, 20, 40, 60, 80, 120, 150 and 200 Hz every 1 min. At the end of the contractile protocol, muscle length was measured using a hand-held electronic caliper, fiber bundles removed from the organ bath and trimmed of excess bone and connective tissue, blotted dry and weighed. Muscle weight and Lo were used to calculate normalized forces expressed as N/cm^2^

### ROS measurements

Diaphragm intracellular ROS was measured using 6-carboxy-20,70-dichlorodihydrofluorescein diacetate (DCFH-DA) (Invitrogen, Carlsbad, CA) as previously described^14^ Briefly, diaphragm muscle optimal length was determined as described above followed by incubation with DCFH-DA for 30 min, washed using the physiological saline solution and de-esterified for an additional 30 min at 25°C. All cell-loading and imaging was performed in the dark to prevent light induced oxidation of DCFH-DA. DCF was excited at 470/20 nm using a Sutter Lamda DG-5 Ultra-high-speed wavelength switcher and emission intensity was collected at 535/48 nm on a charge coupled device (CCD) Camera (CoolSNAP MYO, Photometrics, Tucson, AZ) attached to an Axio Observer (Zeiss) inverted microscope (20X objective, 0.5 NA) at a rate of 0.2 Hz. Alterations in the rate of ROS production were baseline corrected and calculated over the final minute of the stretch period.

### Western blotting and Immunoprecipitation

TA muscle from hind limb of the mice were isolated and lysed with ice-cold RIPA buffer containing protease and phosphatase inhibitors, and centrifuged at 13,500rpm for 10 min. The supernatant was retained, aliquoted, and the protein content was quantified using the bicinchoninic acid (BCA) Assay Kit. A volume corresponding to 10 μg of protein was diluted with a Laemmli sample buffer (BioRad), and heated at 100 °C for 5 min. Protein samples were separated via 4- 15% and 4-20% SDS-PAGE and then transferred to polyvinylidene difluoride (PVDF) membranes, which were blocked with 5% non-fat milk or 5% BSA (dissolved in Tris-buffered saline, pH 7.4, and 0.2% Tween 20, TBST) for 60 min at room temperature.

For immunoprecipitation, total protein lysates were prepared and quantified as described above. Protein lysates (1000μg) were incubated with protein A magnetic beads (BioRad) as per the manufacturer’s instructions. Protein complexes were analyzed by western blotting using indicated antibodies (Beclin-1, ATG14L, Bcl-2, JIP-1, JNK, KIF5B, KLC, SNAP29, VAMP8, STX17, Prx2, acetyl lysine). Immunoblots were incubated overnight at 4 °C with primary antibodies. The following day, membranes were washed and incubated with secondary antibodies for 60min at room temperature. Clarity western ECL substrate (Bio-Rad) was added on the membrane and the signals were visualized by ChemiDoc Imaging system (Bio-Rad). The bands were quantified using densitometric analysis by the ImageLab Software.

### Histology and Immunofluorescence

The skeletal muscles (TA and GAS) were carefully excised from hind limb of mice and post-fixed in 10%formalin overnight. Paraffin blocks were then prepared after dehydration, clearing, and wax impregnation. Transverse Sections (T.S.) of 4μm thickness were cut with a rotary microtome, deparaffinized in xylene and gradient of alcohol were subjected to histological or immunofluorescence staining.

For H&E staining, the slides were stained in hematoxylin solution for 20–30 min and rinsed in running water three times. Slides were then stained in eosin solution for 1–2 min, dehydrated in graded ethanol and xylene, and covered using DPX paramount for further imaging. For immunofluorescence, serial muscle cross-section (4μm) were cut from each paraffin block by rotatory microtome. The sections were placed on coated slides (Denville Scientific Inc.) and dried at 60 °C in a hot air oven. Sections were deparaffinized, rehydrated, and underwent an antigen retrieval method using citrate-EDTA solution incubated at 96 °C for 20 min in water bath. After dipping the slide in distilled water, the sections were blocked with 3% BSA for 60min at room temperature, and incubated with the primary antibodies overnight at 4°C followed by secondary antibodies at room temperature for 1 h. Primary antibodies include SNAP29, VAMP8, CD68, and α-laminin. Secondary antibodies include Alexa Fluor 488 and 594. Tissue sections were mounted with Prolonged Gold Antifade with DAPI (Invitrogen). All images were taken with Fluorescent microscope (ECHO, CA) and analysis was carried out by ImageJ software.

For CD68 quantification, images acquired from five different optical fields of GAS muscle cross-section (40X magnification) and the number of CD68+ immune cells were counted per area (μm^2^).

The CSA and centralized nuclei were measured based on α-laminin staining and DAPI stained nuclei respectively. CSA was quantified by minimum Feret’s diameter and central nucleation was quantified as % fibers with central nuclei. TA muscle cross-sections were used for quantification. All the immunofluorescence images were quantified using Image J software (NIH, USA).

### Colocalization analysis

For double immunofluorescence, two primary antibodies (SNAP29 and VAMP8) were incubated on the same gastrocnemius (GAS) tissue sections. To quantify the degree of colocalization between the anti- SNAP29 and anti-VAMP8 staining, Pearson’s correlation coefficient was performed using ImageJ Software.

### RNA extraction and Gene expression analysis

Total RNAs were extracted from gastrocnemius (GAS) muscles using TRIzol reagent (Invitrogen) according to the manufacturer’s protocol. Gene expression was verified by using cDNA synthesis kit (Company name) for reverse transcription and Real Mastermix for PCR following manufacturer’s instructions. The primers used were as follows:

BECN1-FWD (5’ AGG CTG AGG CGG AGA GAT T-3’), BECN1-REV (5’- TCC ACA CTC TTG AGT TCG TCA T-3’); ATG5-FWD (5’-ATCAGACCACGACGGAGCGG-3’),

ATG5-REV (5’-GGCGACTGCGGAAGGACAGA-3’); ATG12-FWD (5’- ACAAAGAAATGGGCTGTGGAGC-3’),

ATG12-REV (5’- GCAGTAATGCAGGACCAGTTTACC-3’);

ATG14L-FWD (5’- GCAGCTCGTCAACATTGTGT-3’), ATG14L-REV (5’-TGCGTTCAGTTTCCTCACTG-3’); VPS34-FWD(5’-TGTCAGATGAGGAGGCTGTG-3’), VPS34-REV(5’-CCAGGCACGACGTAACTTCT-3’);

GABARAPL1-FWD (5’- CGGTCATCGTGGAGAAGGCT-3’),

GABARAPL1-REV (5’- CCAGAACAGTATAACGGCAACTCC-3’); FIP200-FWD(5’-ACCGTGCACCTGCTATTCCT-3’),

FIP200-REV (5’ CATCATGGACAAGCCCTTCA-3’);

UVRAG-FWD (5’ CAA GCT GAC AGA AAA GGA GCG AG-3’), UVRAG-REV (5’- GGA AGA GTT TGC CTC AAG TCT GG-3’); HDAC6-FWD(5’-AAGTGGAAGAAGCCGTGCTA-3’),

HDAC6-REV (5’- CTCCAGGTGACACATGATGC -3’)’; MEC17/ATAT1-FWD (5’- ACTGA AGGAC ACCTC AGCCC GA -3’), MEC17/ATAT1-REV(5’- TACCT CATTG TGAGC CTCCC GG-3’)

### Evan’s Blue Dye uptake

To assess the membrane integrity of WT*, mdx*, and TubA treated *mdx* mice, Evans Blue Dye (EBD) was injected to the animal as described by Millay et al ^67^. EBD (10mg/ml) was dissolved in PBS (pH 7.4) and sterilized by passage through membrane filters with a 0.2-mm pore size. Mice were injected intraperitoneal *(i.p.),* with 0.1 ml/10g body weight with EBD 24h prior to the completion of two week treatment with TubA. After 24h, DIA muscles were harvested from all the mice groups and sectioned using rotatory microtome. EBD positive fibers showed a bright red emission under fluorescent microscope. All sections were examined, photographed and muscle fiber counted under a fluorescence microscope (ECHO, CA).

### TUNEL ASSAY

GAS isolated from hind limb of mice fixed in 4% formalin overnight and then paraffin-embedded muscles sections (4μm) were assessed by terminal deoxynucleotidyltransferase-mediated dUTP nick end labeling (TUNEL) assay with an in situ Cell Death Detection Kit (Biovision Inc., CA, USA) as per the manufacturer’s instructions. Antigen retrieval was performed and slides were then blocked in 3% bovine serum albumin for 1hr at RT and incubated in anti-α-laminin overnight at 4°C. The following day, slides were incubated in donkey anti-rabbit AF594 and then stained with Prolong gold antifade reagent with DAPI (Invitrogen).Tissue sections were examined under fluorescence microscope (ECHO, CA). TUNEL positive nuclei were counted manually as a co- localization of TUNEL + nuclei with DAPI and expressed as a percentage of the TUNEL + nuclei per total number of DAPI stained nuclei obtained from five different optical field per muscle section (40X Magnification). Total number of DAPI stained nuclei were counted using Image J software (NIH).

### Statistical analysis

Statistical analysis were performed using Origin Pro Software (OriginLab Corporation, Northhampton, MA, USA). Differences between two groups were analyzed using Student’s *t*-tests and normality tests. Differences between means of multiple groups were analyzed by one or two-way analysis of variance *ANOVA* followed with Tukey *post-hoc* tests. Values of *p*<0.05 (95% confidence) were considered statistically significant. Data are represented by box plots with mean (empty checker box within the box), median (solid bar), SD (whisker), SE (box). The number of mice used in groups for each parameter are indicated in figure legends. Recovery index (RITubA) was calculated as ((*mdx*+TubA)-(*mdx*))/((WT)-(*mdx*))x100 ^66^.

## Acknowledgement

Research reported in this publication was supported by the National Institute of Arthritis and Musculoskeletal and Skin Diseases of the National Institutes of Health under Award Number RO1AR061370 and a Mrs. Clifford Elder White Graham Endowed Research Fund Awarded to G.G.R.

## Author Contributions

A.A. and G.G.R conceived and design the experiments; A.A. performed the experiments, analyzed data and wrote the manuscript; E.L.C. and G.G.R. performed ex-vivo muscle contractility experiments; C.L.C. performed stretch induce ROS experiment; B.A.C processed co-localization data. G.G.R. edited the final manuscript. All authors reviewed, edited and approved the final version of manuscript.

## Material and Correspondence

Correspondence to George G Rodney

---

**Fig.S1.**
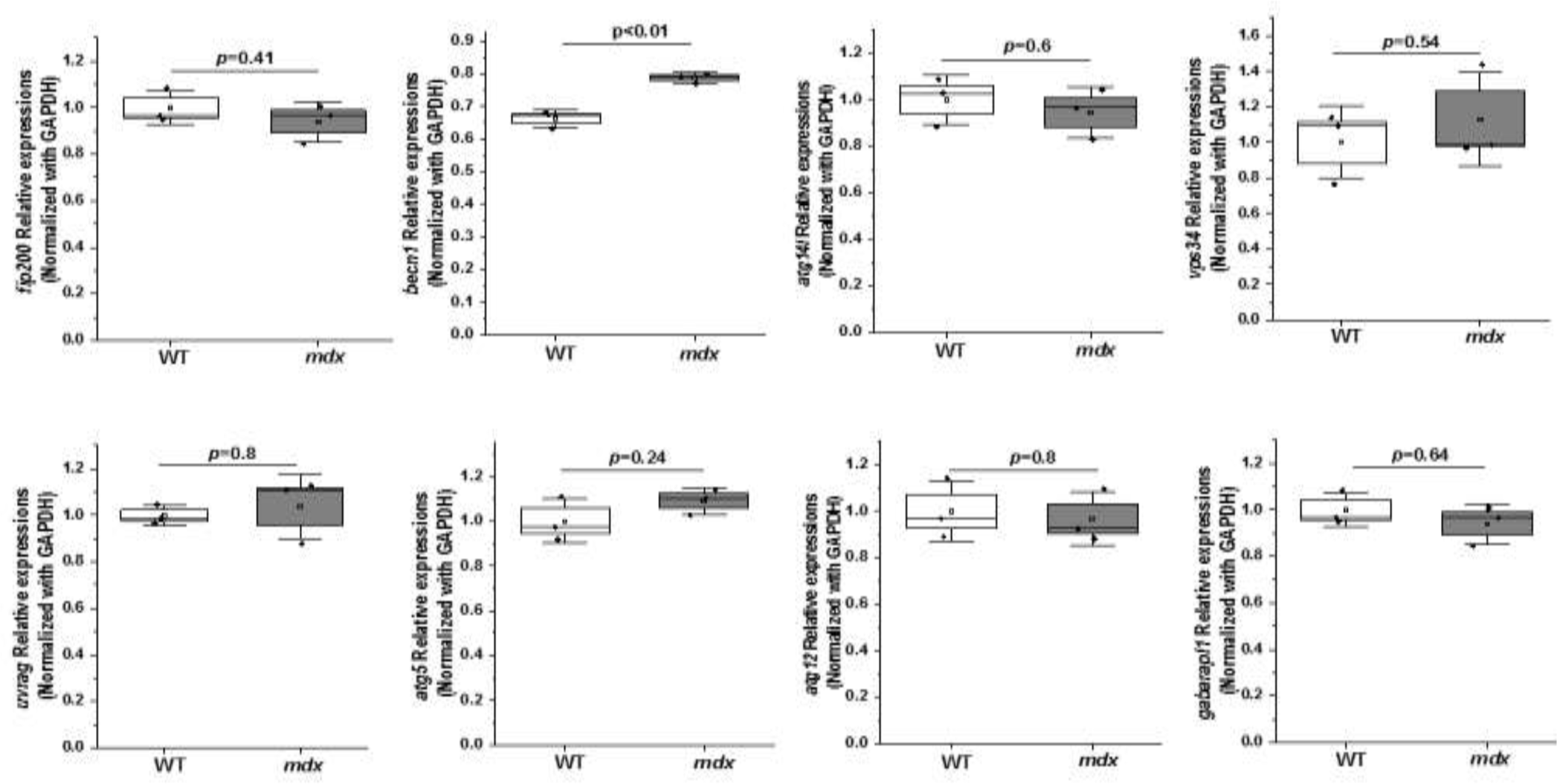
Gene expressional analysis of autophagy regulatory markers by Reverse-transcriptase PCR. Graph represents relative gene expressions of *fip200, becn1, atg14l, vps34, uvrag, atg5, atg12, gabarapl1*normalized with GAPDH (n=3 per group). Statistical difference between two groups were determined using two-sample student-*t-*test.

**Fig.S2.**
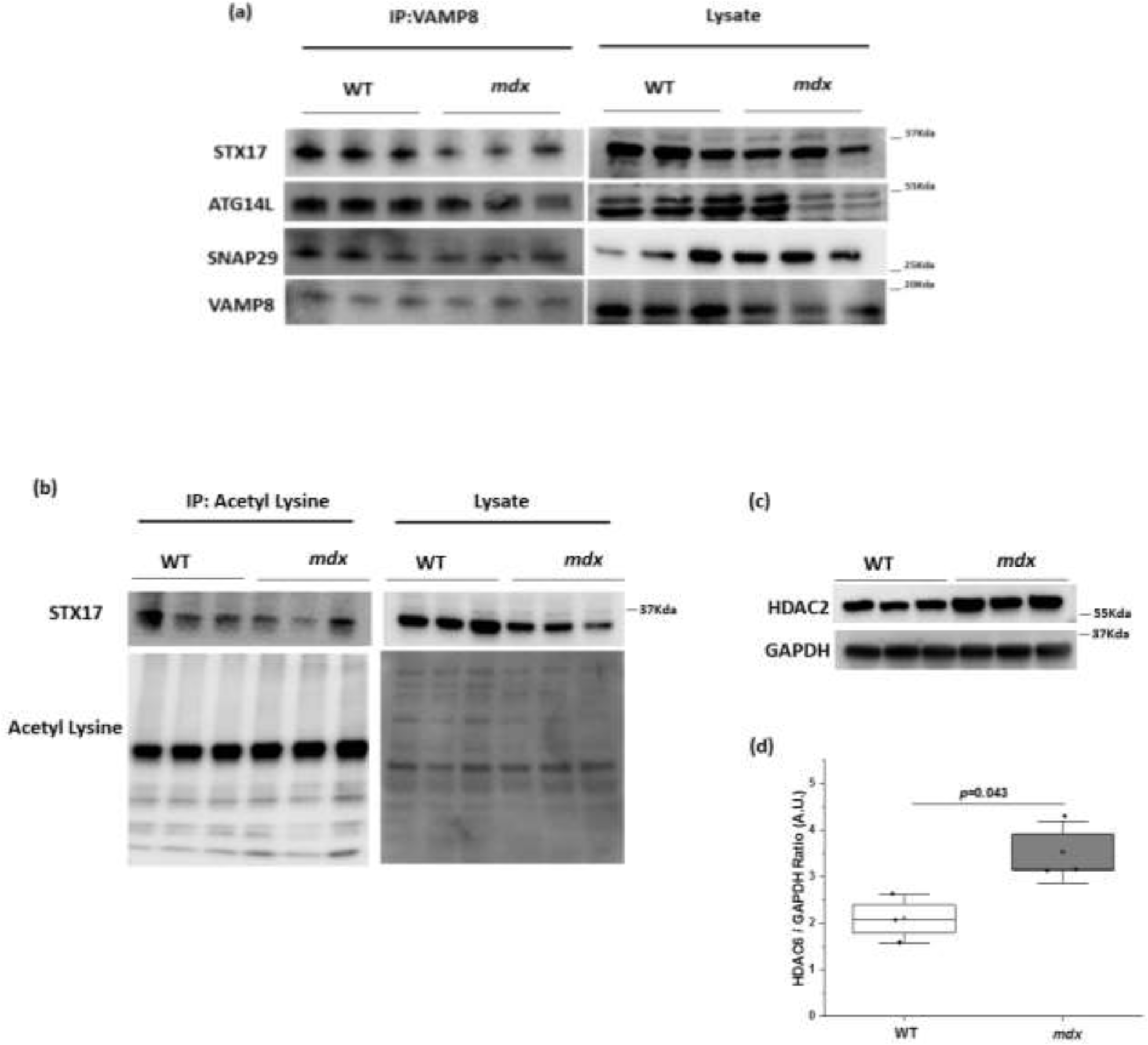
Impaired autophagosome-lysosome fusion in *mdx* mice. **(a)** Skeletal muscle lysate was prepared and subjected to immunoprecipitation using anti-VAMP8, and the associated STX17, ATG14L, and SNAP29 were determined using immunoblotting (b) Anti-acetyl-lysine was immunoprecipitated, and bound STX17 was detected by western blotting using anti-STX17 antibody (c-d) Immunoblot represents protein expressions of HDAC2. GAPDH served as a loading control. Densitometry analysis of immunoblot represented by graph. (n=3 per group in all Figure panels). Statistical difference between two groups were determined using two-sample student-*t-*test.

**Fig.S3.**
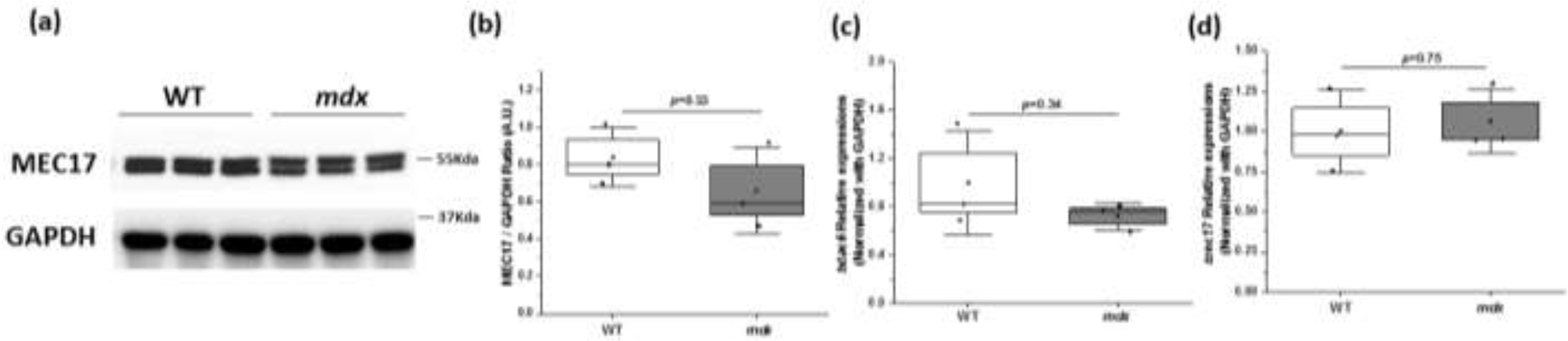
Expressional analysis of acetylase and deacetylase enzymes in *mdx* mice. **(a-b)** Immunoblot represents protein expressions of MEC17. GAPDH served as a loading control. Densitometry analysis of immunoblot represented by graph **(c-d)** Graph represents relative gene expressions of deacetylase enzyme (*hdac6*) and acetylase enzyme (*mec17*) normalized with GAPDH. (n=3 per group in all Figure panels). Statistical difference between two groups were determined using two-sample student-*t-*test.

**Fig.S4.**
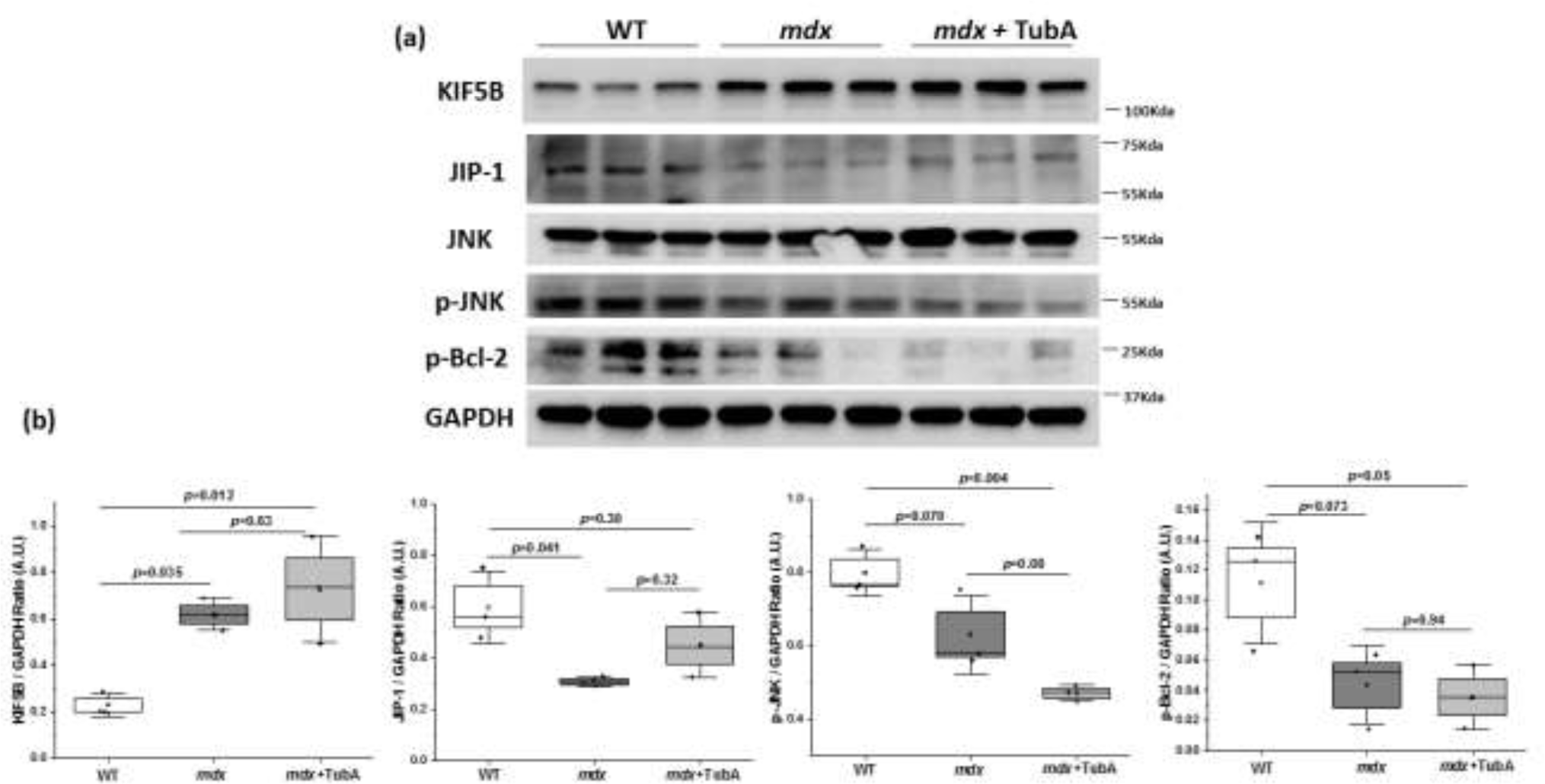
Effect of HDAC6 inhibition on Kinesin/JIP/JNK pathway in *mdx* mice. **(a-b)** Protein expressions of Kinesin/JIP/JNK complex in WT, *mdx* and TubA treated *mdx* mice (n=3 per group). GAPDH served as a loading control. Densitometry analysis of immunoblot represented by graph. Statistical difference between groups were determined using ANOVA with Tukey’*s post hoc* test.

**Fig.S5.**
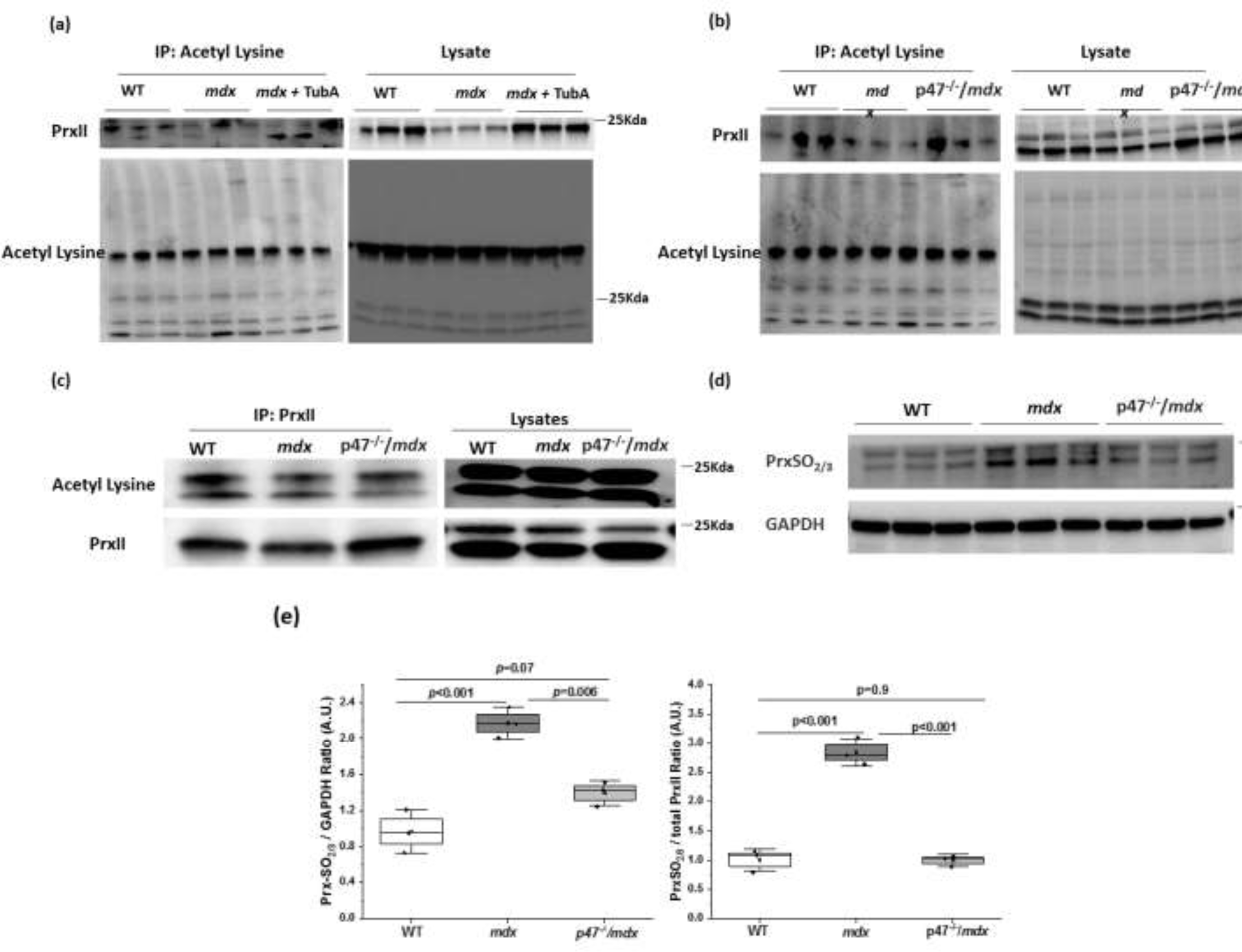
Acetylation status of PrxII in TubA treated *mdx* and genetically ablated Nox-2 *mdx* mice. **(a)** Skeletal muscle lysate was prepared and subjected to immunoprecipitation using anti-acetyl-lysine, and the acetylation levels of PrxII was determined with the anti-PrxII antibodies in WT, *mdx*, and TubA treated *mdx* mice (n=3 per group). Input blot of PrxII for WT *mdx* and *mdx*+TubA group in Fig.S5a are the same as shown in **Fig.6b (b)** Immunoprecipitation was performed using anti-acetyl-lysine, and the acetylation levels of PrxII was determined with the anti-PrxII antibodies in WT, *mdx*, and p47 ^-/-^/*mdx* mice (n=3 per group) **(c)** Immunoprecipitation was performed using anti-PrxII, and the acetylation levels of PrxII was determined with the anti-acetyl lysine antibodies in WT, *mdx*, and p47 ^-/-^/*mdx* mice **(d-e)** Immunoblot represents hyperoxidation state of PrxII (PrxSO2/3) in WT, *mdx*, and p47 ^-/-^/*mdx* mice (n=3 per group). Densitometry analysis of western blots and the ratio of PrxSO2/3 and total PrxII were represented by graph. GAPDH served as a loading control. Statistical difference between groups were determined using ANOVA with Tukey’*s post hoc* test.

**Fig.S6.**
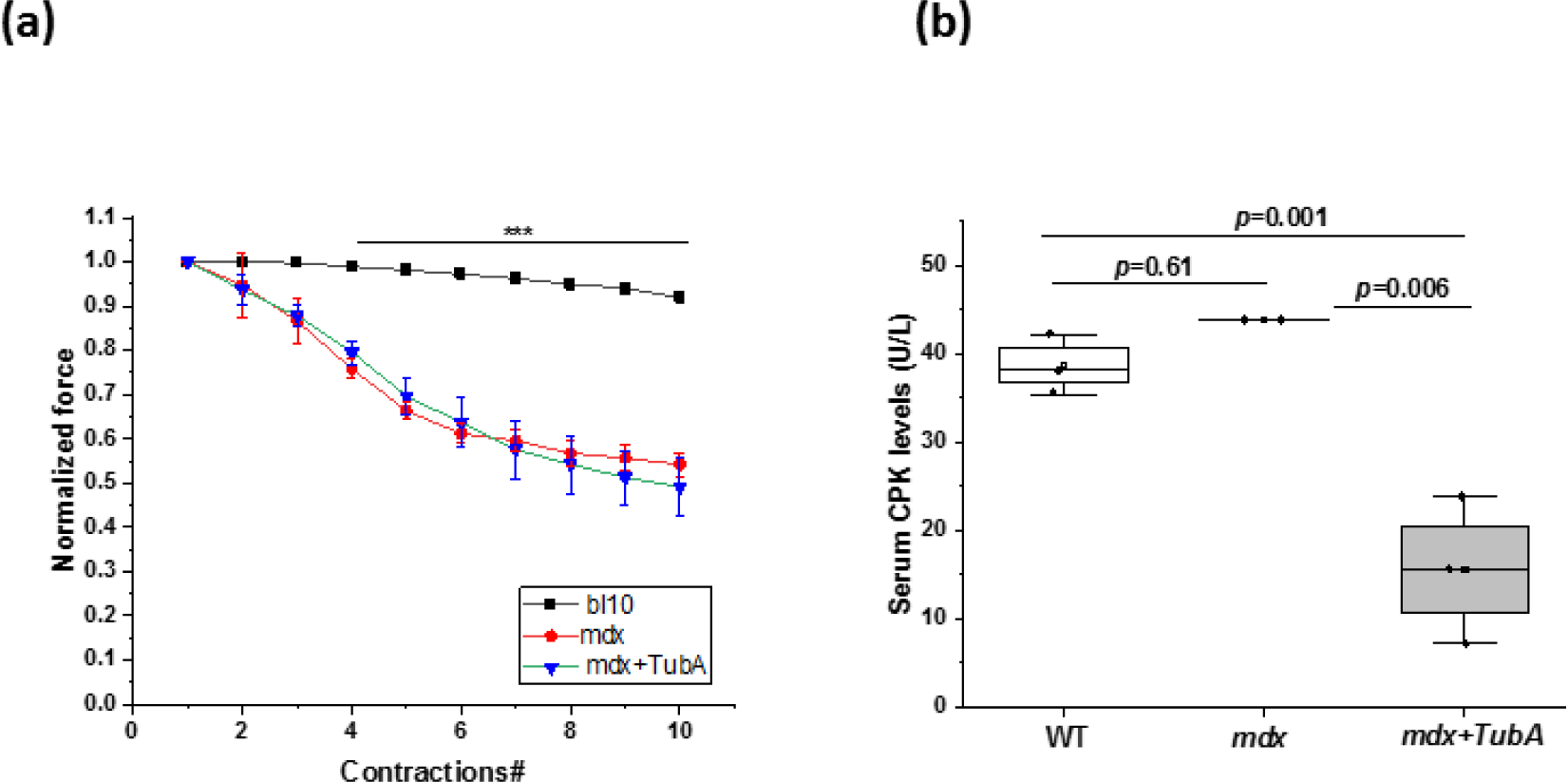
Effect of TubA on muscle contraction and damage at different dose and time point. **(a)** HDAC6 specific inhibitor, TubA at 30mg/kg b.w. injected into mdx mice intraperitoneally for 4 weeks every other day. After completion, EDL muscle isolated from WT (n=2, *mdx (n=3)*, TubA treated *mdx (n=3)* mice and Eccentric contraction induced force drop (normalized to the first contraction) was performed**. (b)** Serum creatine kinase quantitate by ELISA from WT (n=3), *mdx (n=2)*, TubA treated *mdx (n=3)* mice. Statistical difference between groups were determined using ANOVA with Tukey’*s post hoc* test.***p<0.001 vs *mdx* and *mdx* +TubA for **panel a**.

**Supplementary Table 1.**
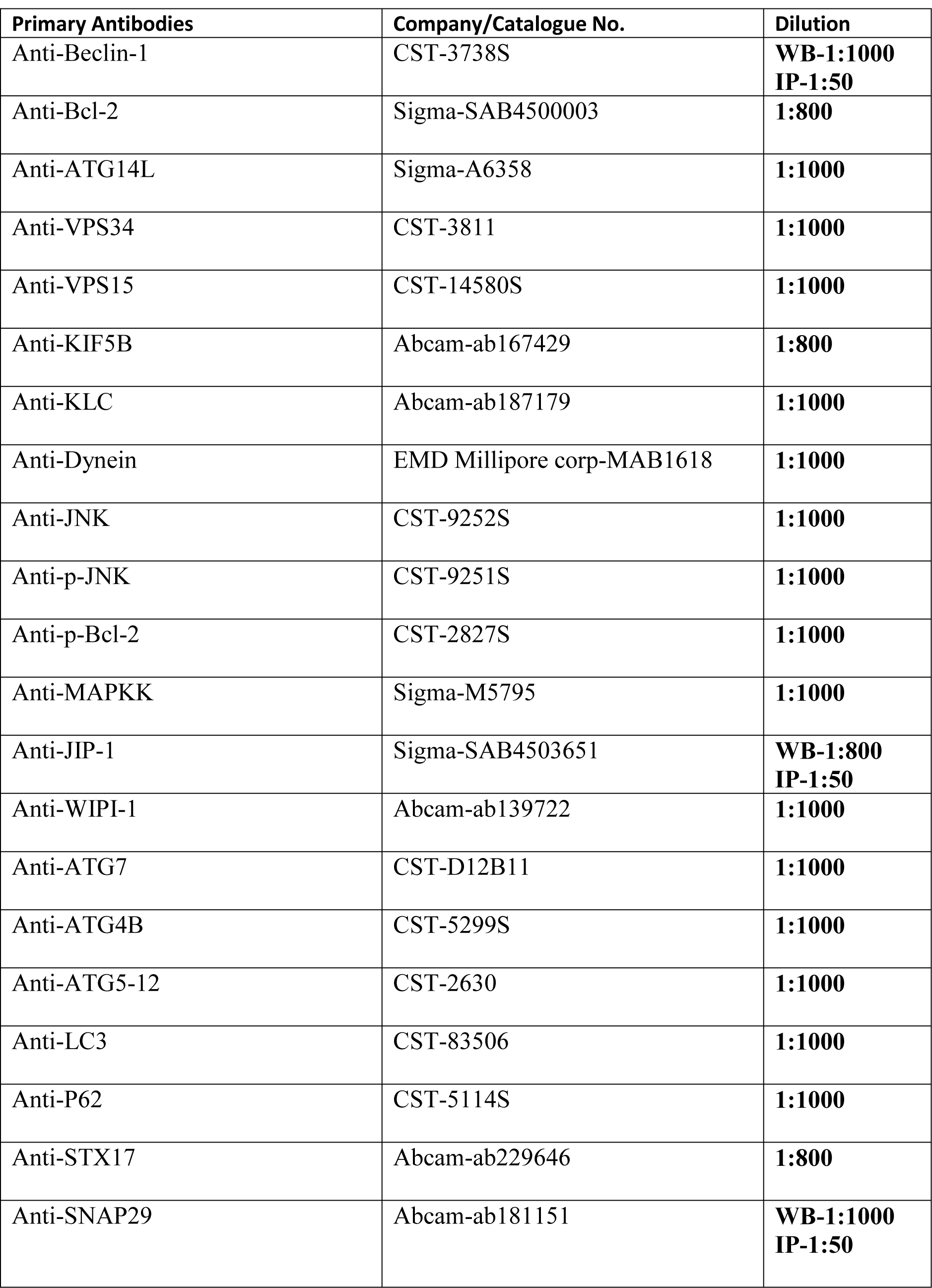
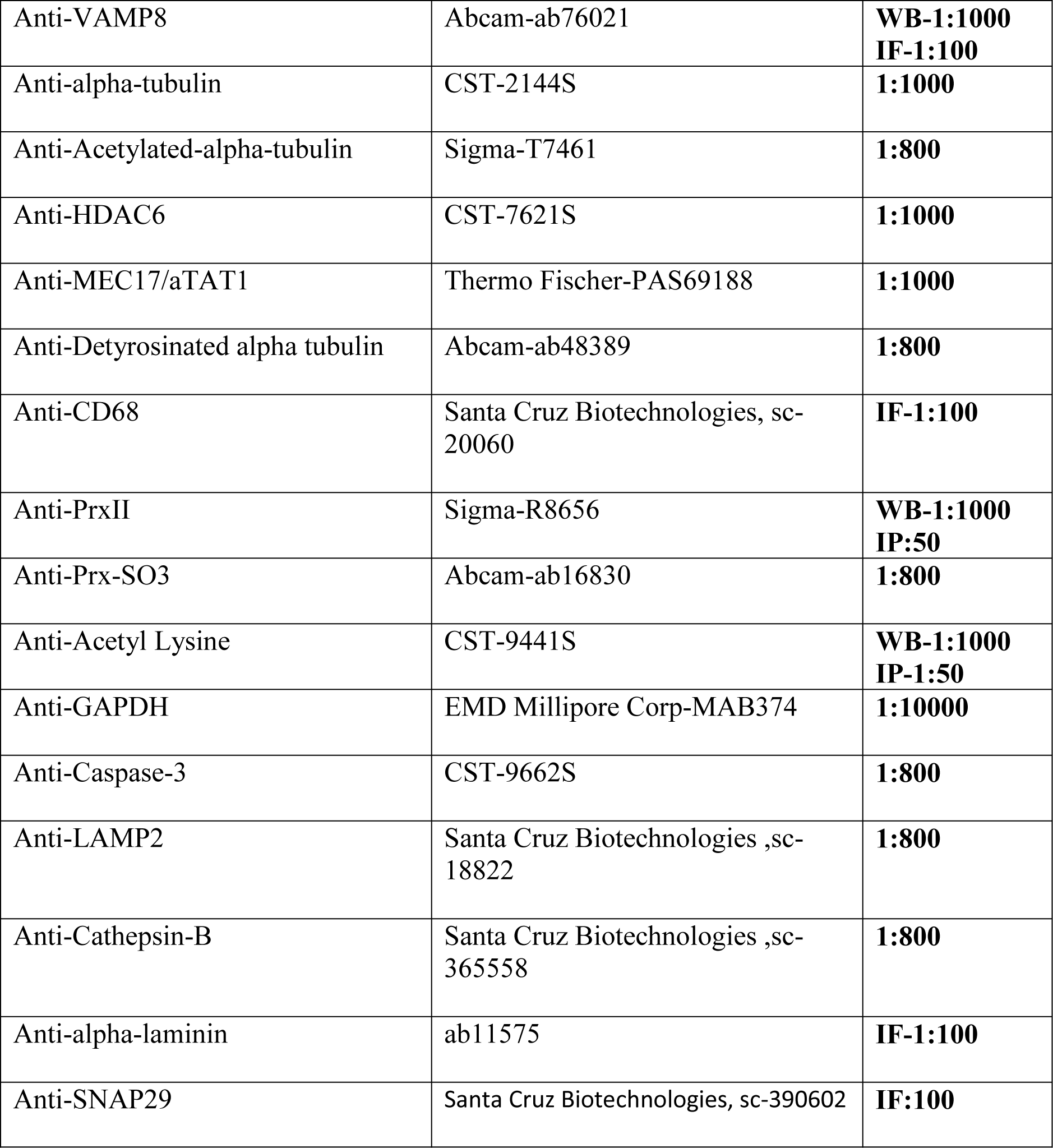
Primary Antibody dilutions

**Supplementary Table 2.** Secondary Antibodies

| **Secondary antibodies** | **Company/catalogue** | **Dilution** |
| --- | --- | --- |
| Anti-mouse | Sigma-GENA931 | 1:10000 |
| Anti-rabbit | Sigma-GENA934 | 1:10000 |
| Alexa-fluor 594 donkey anti-rabbit | Invitrogen A21207 | 1:1000 |
| Alexa –Fluor-488-donkey anti-rabbit | Invitrogen A21206 | 1:1000 |
| Alexa-Fluor-594-goat anti-mouse | Invitrogen A11032 | 1:1000 |

